# Fluorescence hybridization chain reaction enables localization of multiple molecular classes combined with plant cell ultrastructure

**DOI:** 10.1101/2024.01.29.577761

**Authors:** Yunqing Yu, David Huss, Mao Li, Janithri S. Wickramanayake, Sébastien Bélanger, Anastasiya Klebanovych, Blake Meyers, Elizabeth A. Kellogg, Kirk J. Czymmek

## Abstract

**Background:** Recent developments in hybridization chain reaction (HCR) have enabled robust simultaneous localization of multiple mRNA transcripts using fluorescence *in situ* hybridization (FISH). Once multiple split initiator oligonucleotide probes bind their target mRNA, HCR uses DNA base-pairing of fluorophore-labeled hairpin sets to self-assemble into large polymers, amplifying the fluorescence signal and reducing non-specific background. Few studies have applied HCR in plants, despite its demonstrated utility in whole mount animal tissues and cell culture. Our aim was to optimize this technique for sectioned plant tissues embedded with paraffin and methacrylate resins, and to test its utility in combination with immunolocalization and subsequent correlation with cell ultrastructure using scanning electron microscopy.

**Results:** Application of HCR to 10 µm paraffin sections of 17-day-old *Setaria viridis* (green millet) inflorescences using confocal microscopy revealed that the transcripts of the transcription factor *KNOTTED 1* (*KN1*) were localized to developing floret meristem and vascular tissue while *SHATTERING 1* (*SH1*) and *MYB26* transcripts were co-localized to the breakpoint below the floral structures (the abscission zone). We also used methacrylate de-embedment with 1.5 µm and 0.5 µm sections of 3-day-old *Arabidopsis thaliana* seedlings to show tissue specific *CHLOROPHYLL BINDING FACTOR a/b* (*CAB1*) mRNA highly expressed in photosynthetic tissues and *ELONGATION FACTOR 1 ALPHA* (*EF1*α**) highly expressed in meristematic tissues of the shoot apex. The housekeeping gene *ACTIN7* (*ACT7*) mRNA was more uniformly distributed with reduced signals using lattice structured-illumination microscopy. HCR using 1.5 µm methacrylate sections was followed by backscattered imaging and scanning electron microscopy thus demonstrating the feasibility of correlating fluorescent localization with ultrastructure.

**Conclusion:** HCR was successfully adapted for use with both paraffin and methacrylate de-embedment on diverse plant tissues in two model organisms, allowing for concurrent cellular and subcellular localization of multiple mRNAs, antibodies and other affinity probe classes. The mild hybridization conditions used in HCR made it highly amenable to observe immunofluorescence in the same section. De-embedded semi-thin methacrylate sections with HCR were compatible with correlative electron microscopy approaches. Our protocol provides numerous practical tips for successful HCR and affinity probe labeling in electron microscopy-compatible, sectioned plant material.

## BACKGROUND

Characterization of RNA molecules and their downstream products at the cellular and subcellular level is essential to understand gene function and transcript regulation within the complex cellular environment of multicellular organisms including plants. Spatiotemporal expression patterns of genes can be detected using promoter-driven reporter genes expressed in transgenic plants, or *in situ* hybridization (ISH). The latter is particularly useful in species where generation of transgenic plants is unavailable, time-consuming or costly. Traditional ISH uses long (150-1500 bases) anti-sense, single-stranded RNA riboprobes tagged with a hapten such as digoxygenin (DIG), biotin, or fluorescein with uracil (1–3).The haptens are subsequently detected by specific antibodies conjugated with alkaline phosphatase, which catalyzes colorimetric reactions on chromogenic substrates such as nitroblue tetrazolium/ 5-bromo-4-chloro-3-indolyl-phosphate (NBT/BCIP). While this technique has been successfully applied in both whole mount and sectioned tissues in plants (4–9), the diffusion of chromogenic deposition may obscure cellular and subcellular details of signal, and the use of proteases to increase probe permeability may be detrimental to tissue morphology. Additionally, colorimetric ISH does not allow simultaneous detection of multiple genes or other molecules, and therefore is not suitable for studying gene co-expression or the spatial distribution of RNA and other molecules such as proteins and polysaccharides.

Fluorescent ISH (FISH) is an alternative technique that is compatible with multiplexed target detection. FISH relies on fluorophores directly or indirectly conjugated with RNA or DNA probes to visualize signals. Fluorophores can be conjugated to primary antibodies that target RNA probes labeled with a hapten or secondary antibodies and visualized under a fluorescence microscope. However, since this approach lacks an amplification component as in colorimetric ISH, it may not provide sufficient signal-to-noise for visualizing weakly expressed genes. Various techniques have been developed to enhance signal, either through increasing probe numbers, improving detection sensitivity or amplifying the original signal. Single molecule FISH (smFISH) uses a large set of fluorophore-labeled, short, DNA oligonucleotide probes (20 – 22 nucleotides (nt)) dispersed along the target mRNA (10). Super-resolution microscopy can further enhance the detection sensitivity and localization precision of smFISH (11). Direct labeling of DNA probes preserves spatial resolution of the target mRNA compared to methods using enzymes to catalyze chromogenic or fluorogenic reactions. However, direct conjugation of fluorophores to individual probes can be expensive when multiple genes are targeted.

To improve sensitivity, signal amplification technologies have been developed including tyramide signal amplification (TSA) (12), rolling circle amplification (RCA) (9,13), RNAscope (14) and hybridization chain reaction (HCR) (15,16). TSA uses primary antibody conjugated horseradish peroxidase to activate the binding of fluorophore-labeled tyramide with tyrosine residues at or near the peroxidase reaction site (17). TSA has been used to differentiate nuclear and cytoplasmic RNA in Formalin-Fixed, Paraffin Embedded (FFPE) *Arabidopsis* shoot apices (12). Rolling circle amplification (RCA) together with padlock probes and barcoding system allows highly multiplexed *in situ* sequencing by hybridization and has been applied in FFPE maize shoot apex and *Arabidopsis* whole mount root tips to target 90 and 28 genes, respectively (9,13,18,19). The RNAscope® system from ACD Bio (Newark, CA) uses a set of probe pairs, each linked with one half of a Pre-Amp sequence to limit non-specific signal, followed by amplifying the specific fluorescent signal by building a branched DNA molecule (14). RNAscope has been adapted to plant tissues, including FFPE maize and cassava leaf (20,21) and cryo-sectioned barley leaf (22).

Alternatively, HCR FISH adopts an enzyme-free approach and uses two kinetically trapped DNA hairpins linked with a fluorophore. Version 3 enhancements of HCR FISH (23) uses a set of paired 25 nt probes separated by 2 nt, each linked with half of an initiator sequence, resulting in a 45 nt length for each probe. Only when both probes in a pair are fully hybridized to the target, does the full-length initiator sequence become available. This initiates the hybridization and opening of hairpin 1 (H1), which in turn hybridizes and opens hairpin 2 (H2). H2 then unfolds, revealing a complementary strand to H1, and the process is repeated to eventually assemble into a double-stranded polymer of alternated overlapping H1 and H2. Since H1 and H2 are each conjugated to a fluorophore, the result is a large fluorescent polymer tethered to the site of each initial probe pair binding event (Fig. 1). Five unique initiator sequences allow for simultaneous hybridization and detection of multiple target mRNAs (23). HCR FISH has the benefit of being sensitive and quantitative (24). The reagents can be prepared in house or are commercially available through Molecular Instruments (Los Angeles, CA). Recently, HCR FISH has been applied in wholemount tissues of *Arabidopsis* inflorescences and *Arabidopsis*, maize and sorghum roots (13,25,26).

**Figure 1.**
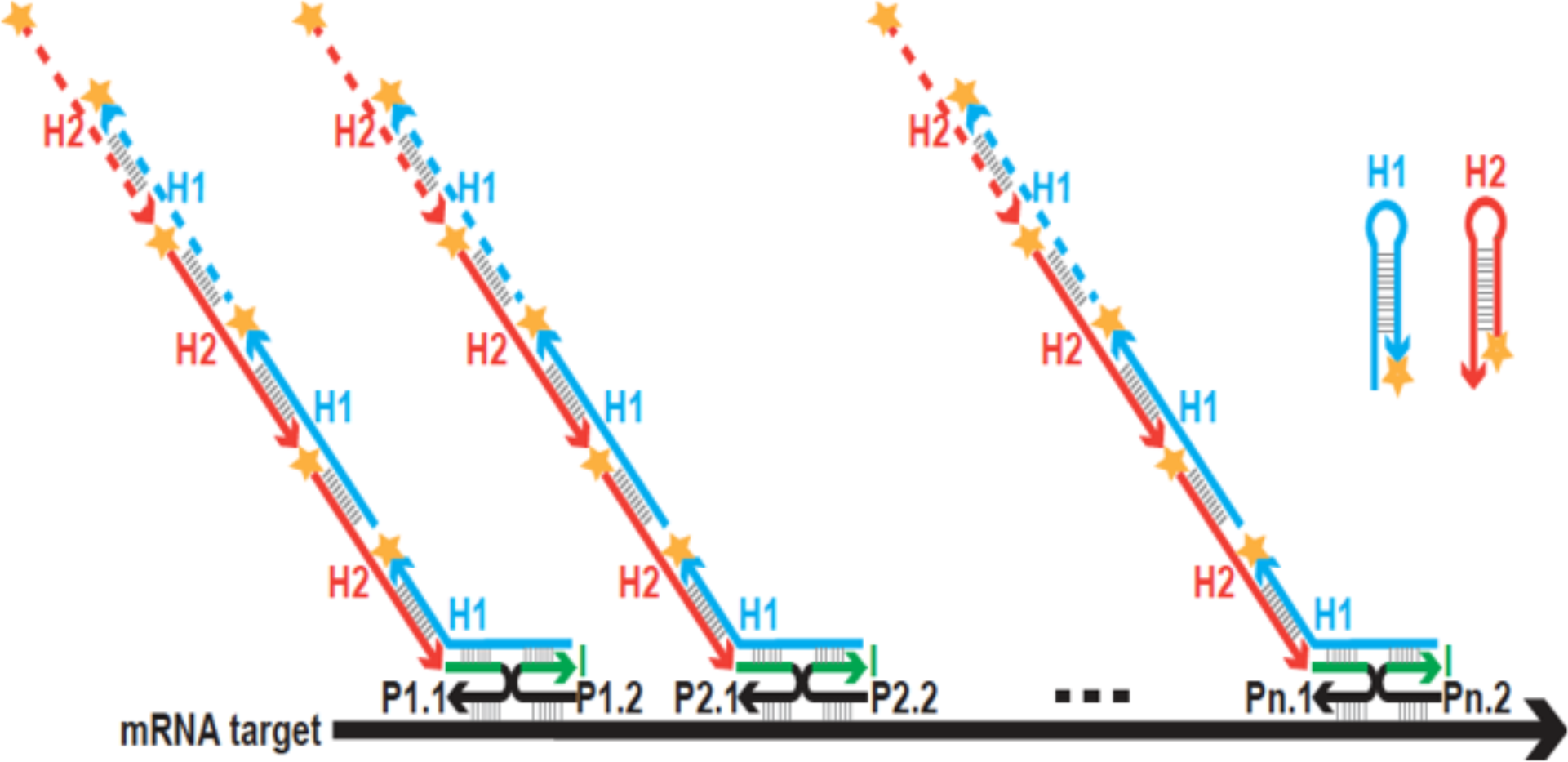
HCR Mechanism. A. The HCR mechanism using split initiator probe pairs. Each anti-sense oligonucleotide probe carries one half of a unique initiator sequence. Once both probes in a pair (P1.1 and P1.2, P2.1 and P2.2, etc.) bind the target mRNA the complete initiator sequence is then available for the fluorescently labeled hairpins (H1 and H2) to self-assemble a tethered polymer. The five unique initiator sequences allow multiple target mRNA’s to be probed and amplified simultaneously utilizing hairpins conjugated to spectrally separated fluorophores.

Thus far, all the above techniques use wholemount or FFPE processed plant tissues. Although subcellular localization has been demonstrated, due to the inherent nature of the typical FFPE fixation and wax-based embedment strategy, in most instances ultrastructural quality is severely compromised or simply not practical. *In situ* hybridization at the transmission electron microscopy level has also been successfully applied in plant sections for abundant mRNA targets and has the benefit of providing sub-cellular ultrastructural context to its distribution (27–29). However, these approaches were limited to a single mRNA target population and combination with immunolocalization proteins was not employed. Methacrylate resins for protein and/or nucleic acids have been successfully applied with improved morphology and subcellular localization compared to FFPE (5,30,31). Furthermore, in this approach, the semi-thin methacrylate sections which are de-embedded with acetone, improved probe accessibility and thus labeling density for light microscopy. Aldehyde fixation and resin de-embedment would be expected to negatively impact morphology compared to osmium tetroxide fixed versus non-de-embedment acrylic and/or epoxy resins. However, this strategy is a recognized compromise to balance adequate probe signal and provide reasonable cellular structure and nevertheless, far superior to paraffin.

Here, we demonstrate the utility of HCR FISH in sectioned FFPE and methyl methacrylate embedded samples. Using an optimized protocol, we show multiplex HCR FISH with the capacity of combining immunofluorescence and other affinity probes with seedlings of *Arabidopsis thaliana* and inflorescence of *Setaria viridis*. Furthermore, for the first time, we correlated HCR FISH with electron microscopy images from the same section, enabling the potential for high spatial resolution of the subcellular localization of mRNA, proteins, and other cell molecules with the long-term goal to localize cellular pathways.

## RESULTS

### HCR detected low-level gene expression in FFPE samples

In order to establish and optimize an HCR FISH protocol for FFPE plant material, we first used panicles from 17-day-old *Setaria viridis*, in which colorimetric ISH was previously demonstrated (12,32). Three transcription factors, *SHATTERING1* (*SH1*, Sevir.9G153200.1), *MYB26* (Sevir.5G293800.1) and *KNOTTED-1* (*KN1*, Sevir.9G107600.1) were used for multiplexed HCR FISH. Both *SH1* and *MYB26* were specifically enriched in the abscission zone below the spikelet based on our previous RNA-Seq analysis (32). However, while *SH1* can be validated by colorimetric ISH, *MYB26* failed to be detected by this method, possibly due to its low absolute expression level (Fig. S1) (32) *KN1* is highly expressed in the apical meristem except the epidermal layer, and the vascular tissues (Fig. S1) (33,34). For each gene, seven to eight pairs of probes spanning the coding sequence (CDS) and 3’ untranslated region (UTR) were designed, and each probe carried half of a unique initiator, specifically, Initiator B2 for *SH1*, B3 for *MYB26* and B4 for *KN1* (Table S1). The initiators can be recognized by specific hairpins conjugated to spectrally separated fluorophores (Alexa Fluor 546 for B2, Alexa Fluor 647 for B3, and Alexa Fluor 488 for B4), allowing simultaneous amplification and detection for all three genes. We observed above background signals of *SH1* and *MYB26* in the abscission zone (AZ), and *KN1* in the floral meristem (Fig. 2A), consistent with the literature (32,33). Negative controls for HCR experiments included samples without probes or hairpins to assess autofluorescence of the tissue (Fig. 2B), and samples with hairpins alone to indicate non-specific amplification (NSA) (Fig. 2C). These controls were essential in determining true signals from background in plant materials, as autofluorescence was observed in our samples most notably with green (488 AF) and red (546 AF) emission channels (Fig. 2B). Our initial experiments using probes without purification identified substantial NSA, especially for the MYB26 probes. Polyacrylamide gel electrophoresis (PAGE) purification of the probes prior to HCR, selecting probe sizes of 45 nt, greatly reduced NSA (Fig. 2C). We therefore used purified probes for subsequent experiments.

**Figure 2.**
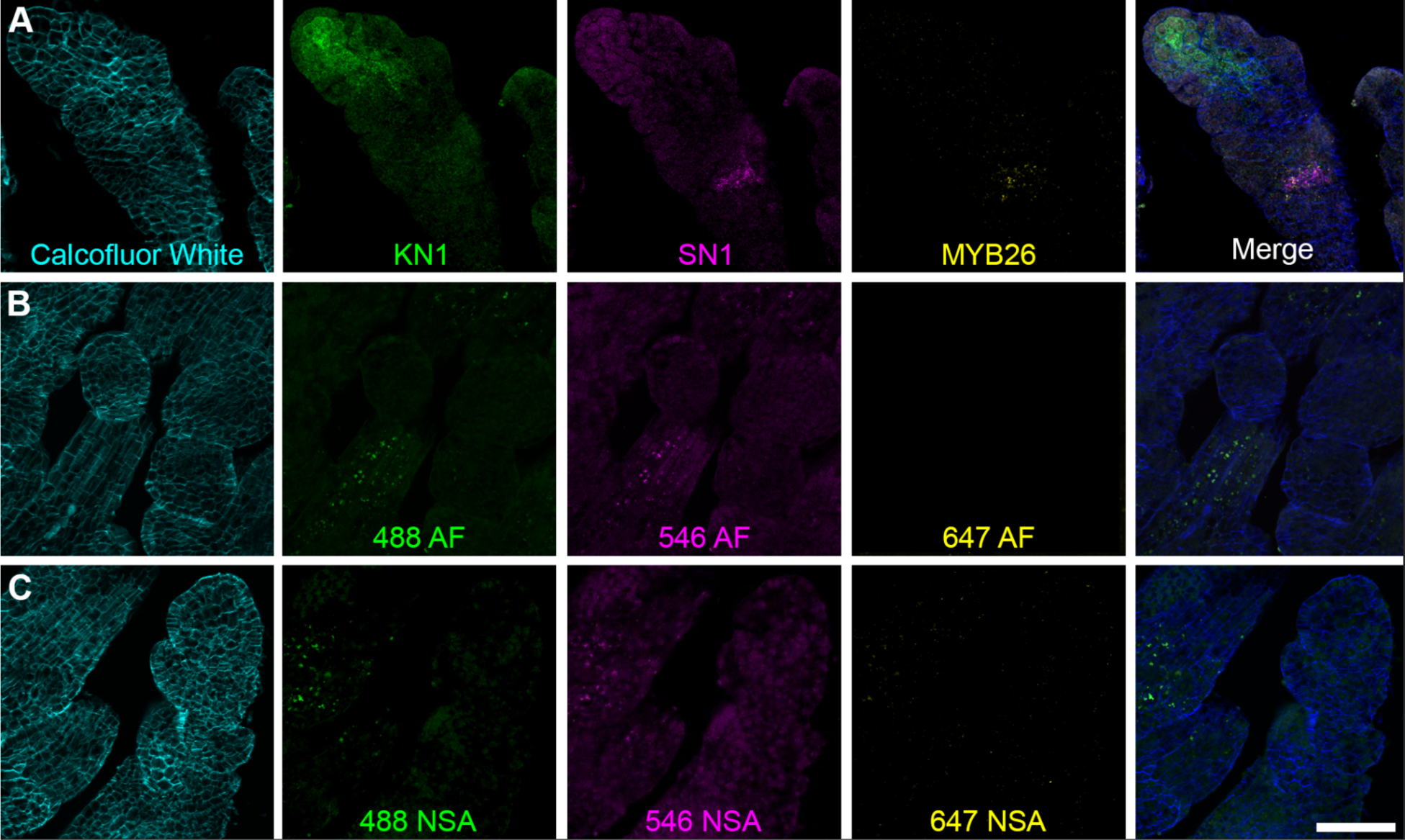
HCR detected multiple mRNA transcripts simultaneously in paraffin sections of *Setaria* inflorescences. A-C. Longitudinal plane through 10 µm paraffin sections of 17-day *Setaria viridis* florets imaged with a 63x/1.2 NA water immersion objective. A single 1.37µm optical slice is shown. A. HCR for KN1, SH1 and MYB26 HCR. KN1: homeotic protein KNOTTED1 (Alexa Fluor 488), SH1: YABBY 2-like protein (Shattering1, Alexa Fluor 546), MYB26: transcription factor MYB86-like (Alexa Fluor 647). B. Autofluorescence (AF) control. No HCR probes or hairpins were used. C. Non-specific amplification control (NSA). Fluorescent hairpins only were used. Cell walls were labeled with Calcofluor White. Scale = 40 µm.

### HCR simultaneously detected mRNA of multiple genes

Next, we tested the capacity of HCR in semi-thin, methyl methacrylate, de-embedded samples. We used 3-day-old *Arabidopsis thaliana* seedlings, in which methacrylate embedded samples were previously demonstrated for colorimetric ISH (5). HCR probe sets were designed against *CHLOROPHYLL BINDING FACTOR a/b* (*CAB1*, At1G29930) expressed in photosynthetic tissues, *ELONGATION FACTOR 1 ALPHA* (*EF1*α**, At1G07940) expressed in actively dividing cells, and *ACTIN 7* (*ACT7*, At5G09810) expressed ubiquitously (Table S1). *CAB1*, *EF1*α** and *ACT7* were probed using B2-Alexa546, B3-Alexa647 and B4-Alexa488, respectively. The formation of hairpin polymers in the presence of a probe pair was confirmed *in vitro* (Fig. S2). HCR applied to 10 µm FFPE sections using confocal microscopy detected signals in the chloroplast-containing cells of cotyledons, shoot apex and hypocotyls for CAB1 and shoot apex for EF1**α** (Fig. 3A and Fig. S3, A and C). However, the *ACT7* signal could not be readily differentiated from the background (data not shown) using our confocal microscope. Semi-thin methacrylate samples (1.5 µm) yielded similar results (Fig. S3, B and D), even for sections at thin as 0.5 µm (Fig. S4), but the morphological properties of the tissues were far superior to the paraffin sections (Fig. 2A, compare Fig. S3 C and D to Fig. S3 A and C). In addition, reduced background signals were observed from these tissues and probes (Fig. 2, B and C).

**Figure 3.**
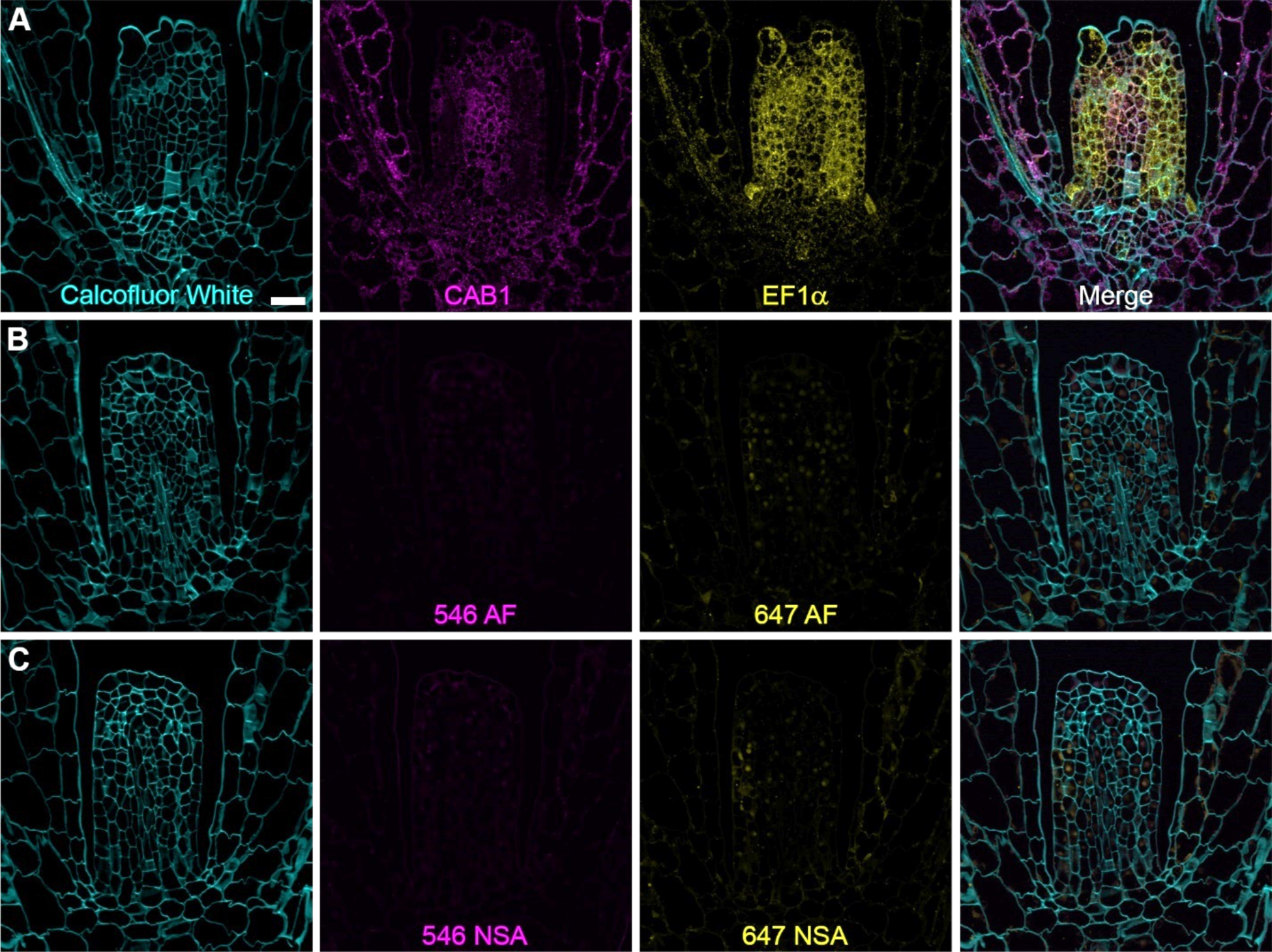
HCR detected multiple mRNA transcripts simultaneously in methacrylate sections of *Arabidopsis* seedling. A-C. Longitudinal plane through 1.5µm methacrylate sections of 3-day *Arabidopsis thaliana* stem apex and cotyledons imaged with a 63x/1.2 NA water immersion objective. A single 1.37µm optical slice is shown. A. HCR for CAB1 (AlexaFluor 546) and EF1*α* (AlexaFluor 647). CAB1: chlorophyll binding protein a/b, EF1*α*: elongation factor 1 alpha. B. Autofluorescence (AF) control. No HCR probes or hairpins were used. C. Non-specific amplification control (NSA). Fluorescent hairpins only were used. Cell walls were labeled with Calcofluor White. Scale = 20µm

### Super-resolution microscopy improved detection sensitivity and resolution of HCR signals

We were unable to detect ACT7 HCR signal in either paraffin or methacrylate embedded sections using our confocal system (Fig 4A). To determine if this was simply due to low-intensity signals, we applied a super-resolution approach with high-sensitivity, complementary metal-oxide-semiconductor (CMOS) cameras and lattice structured illumination microscopy (SIM). In both 10 µm paraffin and 1.5 µm methacrylate embedded sections, we observed individual puncta of Alexa Fluor 488 signal in samples incubated with *ACT7* probes (Fig. 4, B and D) but not with the hairpin control (Fig. 4, C and E), suggesting that the observed signal was not due to autofluorescence or NSA. This also demonstrated the utility of SIM-based super-resolution microscopy in detecting low-intensity HCR signals in plant sections.

**Figure 4.**
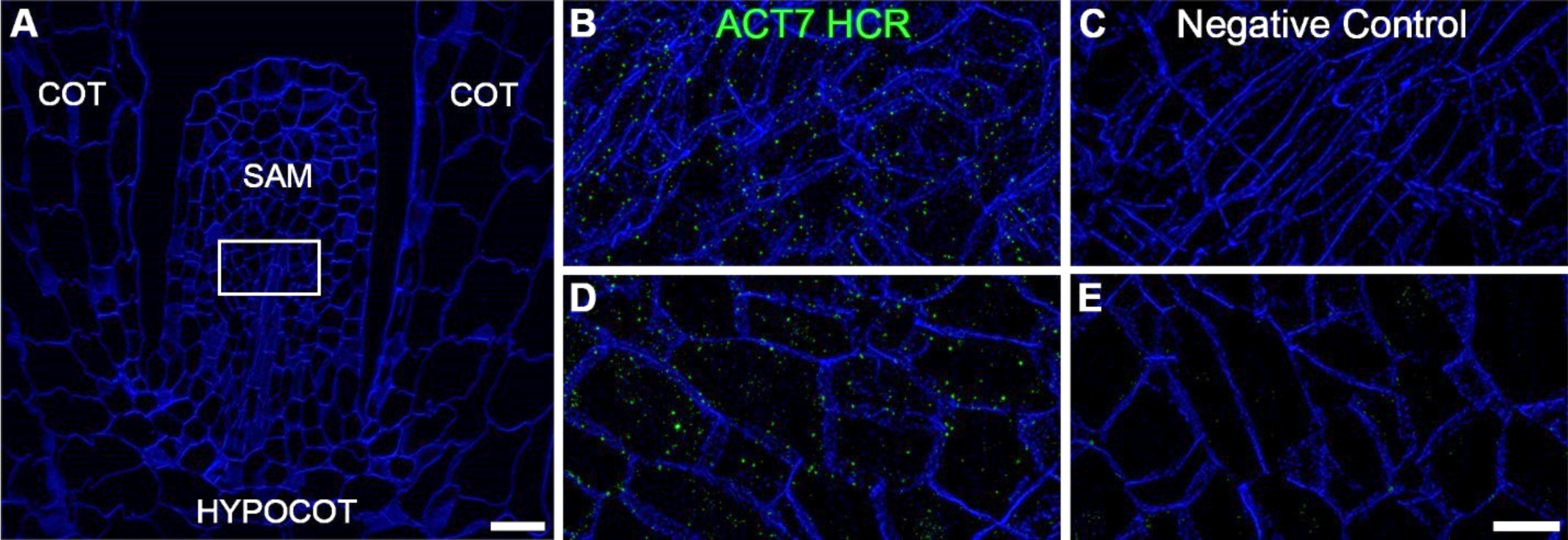
Lattice SIM imaging improved detection sensitivity of low intensity HCR signal. A. A representative confocal microscope image of a 3-day-old *Arabidopsis thaliana* seedling sectioned longitudinally through the shoot apical meristem (SAM), cotyledons (COT) and hypocotyl (HYPOCOT). The bounding box indicates the relative location of B-E. Scale = 20 µm. B. Lattice SIM image of HCR for ACT7 (green) labeled with Alexa Fluor 488 in a 10µm thick paraffin section. C. Alexa Fluor 488 hairpin only control for non-specific amplification (NSA) in a 10 µm paraffin section. D. HCR for ACT7 (green) labeled with Alexa Fluor 488 in a 1.5 µm methacrylate section. E. Green autofluorescence control in a 1.5 µm methacrylate section. Scale = 5 μm. Cell walls stained with Calcofluor White (blue) in all images. HCR and negative control lattice SIM images were taken with identical microscope parameters and displayed with equal pixel brightness settings.

### HCR FISH can be combined with immunofluorescence in both FFPE and methacrylate sections

To test the compatibility of HCR with immunolabeling, we performed a standard immunolocalization protocol after the completion of HCR prior to imaging. With FFPE *Setaria* inflorescences, we observed elevated *SH1* transcript signals restricted in the AZ and the ubiquitous cytoskeletal actin proteins (Fig. 5A), while very low background signals were observed in negative controls with the corresponding hairpins for B2-Alexa546 system and anti-rabbit DyLight650 secondary antibody (Fig. 5B). Likewise, using *Arabidopsis* seedlings embedded in methacrylate, *CAB1* transcripts were readily detected in cotyledons and at low levels in the shoot apex as expected. In the same section, unesterified homogalacturonan, a type of pectic polymer, was detected uniformly in the cell wall using antibody LM19 (Fig. 5C). NSA and non-specific secondary antibody negative controls in sections from the same biological samples demonstrated specificity of our labeled probes (Fig. 5D).

**Figure 5.**
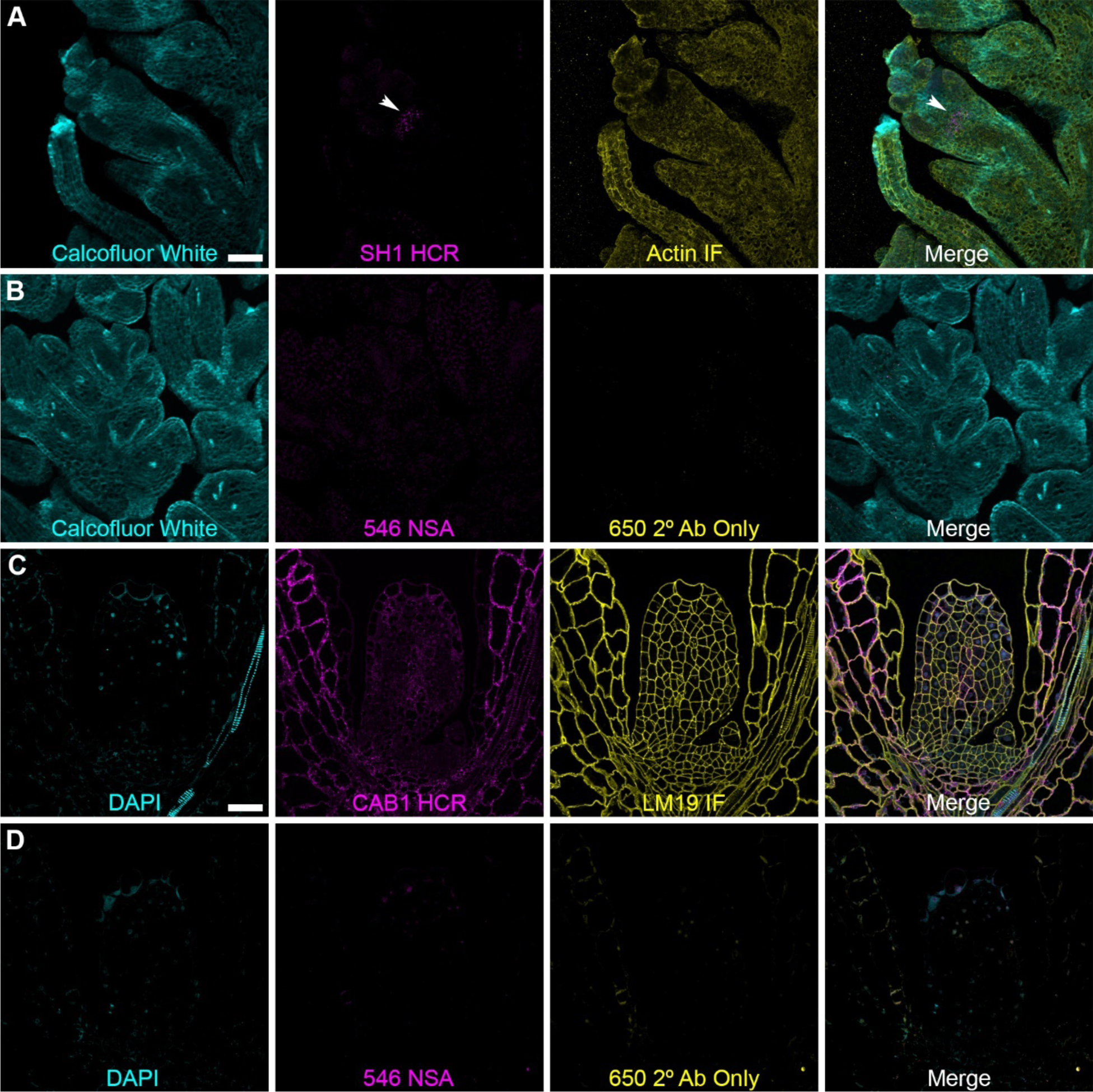
HCR combined with Immunolabeling in the same sections. A. 10 µm paraffin section of the spikelet abscission zone. *SH1*: Shattering 1 mRNA Alexa Fluor 546 (white arrowhead). Actin immunolabeling: Agrisera AS13 2640 Rabbit anti-actin (100 amino acid recombinant peptide actin) detected by Agrisera donkey anti-rabbit IgG DyLight 650. 20x 0.7NA objective, 1.5x zoom, 1.24 AU pinhole = 2.3 µm single optical slice. Scale A & B = 50 µm. B. Non-specific amplification control using only B2 Alexa Fluor 546 hairpins and donkey anti-rabbit IgG DyLight 650 secondary antibodies. Brightness set using the HCR & immunolabeling image then reused on the control images. C. *Arabidopsis* seedling shoot apex in a semi-thin methacrylate section with LM19 immunolabeling and *CAB1* HCR. 63x objective. Scale C & D = 50 µm D. Non-specific amplification control using only B2 Alexa Fluor 546 hairpins and Donkey anti-rat IgG DyLight 650 secondary antibodies. Brightness set using the HCR and immunolabeling image then reused for control images.

### Linear spectral unmixing improved probe signal separation from plant autofluorescence

In some instances, plant cell autofluorescence, which could vary based on tissue and cell type, interfered with our ability to cleanly delineate signal from background, especially when low-intensity signals/expression were observed. To address this, we evaluated the use of spectral confocal microscopy using linear unmixing of reference signals versus the traditional bandpass filter equivalent (559-694 nm) of fluorescence HCR from Alexa Fluor 647 *EF1*α** (compare Figs S5 A and B). It can be readily observed that while some puncta were reliably delineated from cytoplasmic background autofluorescence in the bandpass equivalent (Fig. S5 B blue arrow) a number of puncta overlapped with elevated autofluorescence in chloroplasts, for example (Fig. S5 B white arrows), making definitive localization problematic. The corresponding arrows in the spectrally unmixed data (Fig. S5 A) were readily separated and delineated from autofluorescence background. Furthermore, to better understand the actual spectra in relation to plant autofluorescence with our selected fluorophores and the excitation and emission characteristics of our spectral confocal (ZEISS LSM980 with Quasar detector), we created reference spectra of the pure fluorophores (Calcofluor White and Alexa Fluor 647) from Fig. S5 A & B and autofluorescence used in our test system under identical image acquisition conditions (Figs. S5 C & D). The raw (Fig. S5 C) and normalized (Fig. S5 D) spectra showed the dominant broad spectrum autofluorescence covering a large swath of green and red wavelengths, while tailing off rapidly at shorter blue wavelength and longer red wavelengths. These data emphasized the value of using longer red wavelength fluorophores along with standard bandpass strategies, particularly when low-intensity signals might be anticipated, or in general for best signal-to-noise. We assessed this spectral linear unmixing strategy in methacrylate sections over a large spectral range (411-694 nm) and were able to obtain excellent and clean signal separation with an *A. thaliana* meristem labeled with Calcofluor White (Fig. 6 A), plant autofluorescence (Fig. 6 B), HCR *EF1*α** (Fig. 6 C) and HCR CAB1 (Fig. 6 D) and all probes combined without (Fig. 6 E) and with autofluorescence in the merged images.

**Fig 6.**
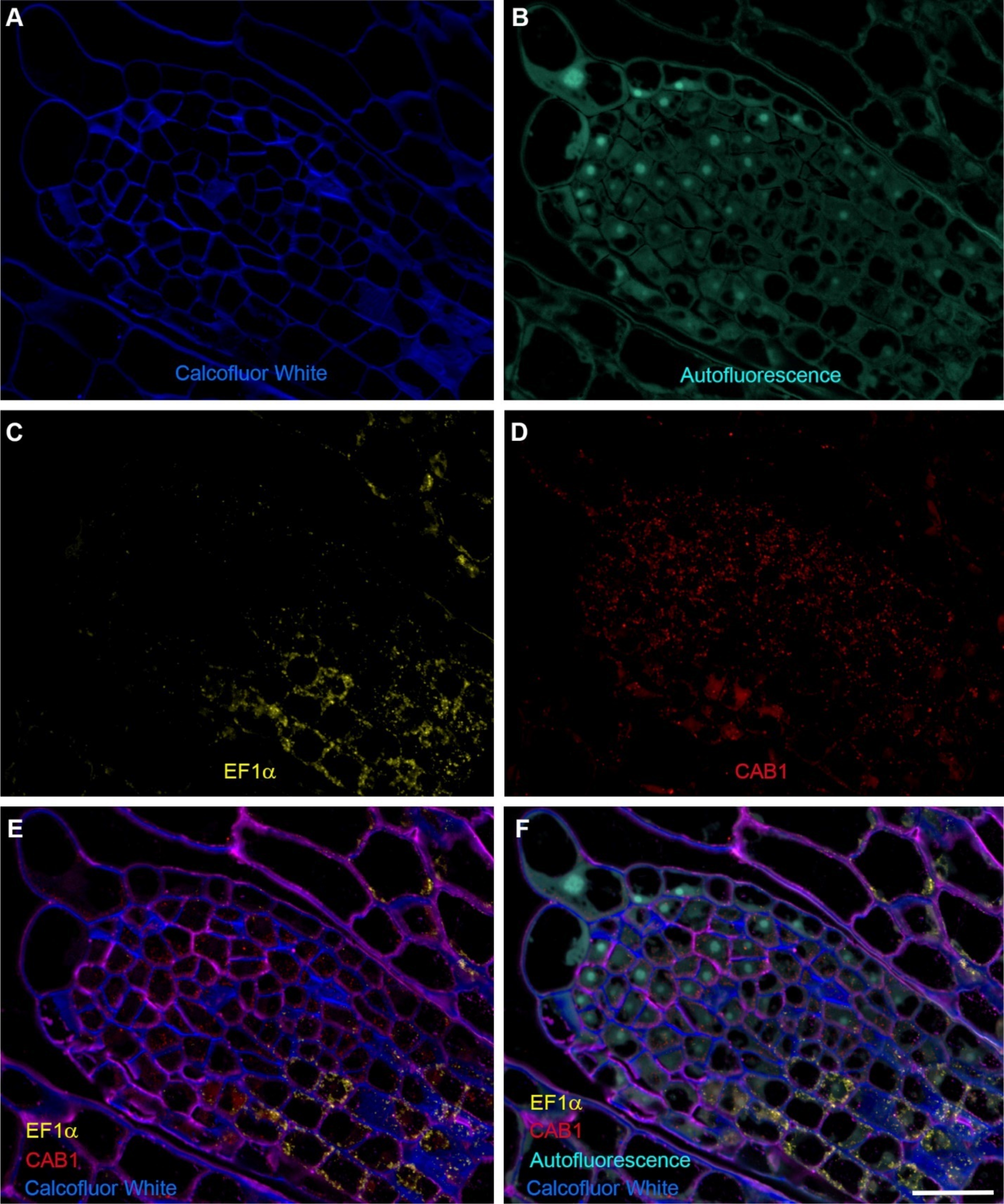
Spectral linear unmixing of methacrylate de-embedded *A. thaliana* meristem showed improved probe separation from plant autofluorescence. A 1.5 µm methacrylate section of an *A. thaliana* meristem labeled with Calcofluor White (A), plant autofluorescence (B), HCR *EF1*α** (C) and HCR CAB1 (D) following linear unmixing using reference spectra for each dye. The resulting clean signals, following linear unmixing, allowed more definitive separation of probes, especially from the substantially cytoplasmic autofluorescence (B). Once achieved, several spectrally distinct fluorescent probes could be combined without (E) or with (F) the autofluorescence signal. Scale = 20 µm.

### Correlative microscopy allowed visualization of cell ultrastructure with HCR FISH

One of our goals was to assess the feasibility of HCR localization (and potentially other probes) with ultrastructure. Since methacrylate de-embedded samples at 1.5 µm significantly improved cellular morphology of the tissues compare to FFPE and allowed for robust localization using fluorescence, we further tested whether the section quality was suitable for ultrastructural observations using backscatter electron detections with field emission scanning electron microscopy (FESEM). In our initial efforts, we found that 1.5 µm sections of the 3-day-old *A. thaliana* meristem at 50 nm x-y pixel resolution allowed the visualization of the cell wall, vacuoles, nucleus, mitochondria, and cytoplasm (Fig. S6 A). Using 50 nm x-y pixel resolution with FESEM was convenient and efficient to perform large area mapping in a reasonable period to assess overall morphology and tissue quality. However, as expected, 5 nm x-y pixel resolution notably improved the clarity of the same sample/location structures (compare Fig. S6 A with B) and more definitive cell details. Nevertheless, with backscatter FESEM, 1.5 µm was too thick and loss of clarity prevented delineation of small organelles or sub-organelle features. We therefore sought to identify a balanced thickness producing sufficient fluorescence signals using fluorescence microscopy and ultrastructural detail. While screening a number of plant tissue types and thicknesses (e.g., Fig. S6), we determined that 0.5 µm methacrylate de-embedded sections provided improvements in clarity as shown in these *A. thaliana* root tip cells at 5 nm pixel resolution (Fig. S6 C and D). Specifically, in the root tissue, the nuclear envelope, endoplasmic reticulum, mitochondria, and cristae were readily identified and resolved. Using the previously established probe set, we successfully detected HCR FISH signals for *EF1*α** in 0.5 µm methacrylate sections of 3-day-old seedling *A. thaliana* cotyledon (Fig. S4). Importantly, following fluorescence imaging, the same sections went through additional osmium tetroxide fixation and heavy metal staining steps to enhance and reveal subcellular structure with FESEM with recognizable nuclei, vacuoles and chloroplasts (Fig. 7). Overlay of the ultrastructure images with HCR FISH found that most of the mRNAs are localized in the cytoplasm, although sporadic HCR puncta were observed in individual nuclei and the cell wall (Fig. 7).

**Figure 7.**
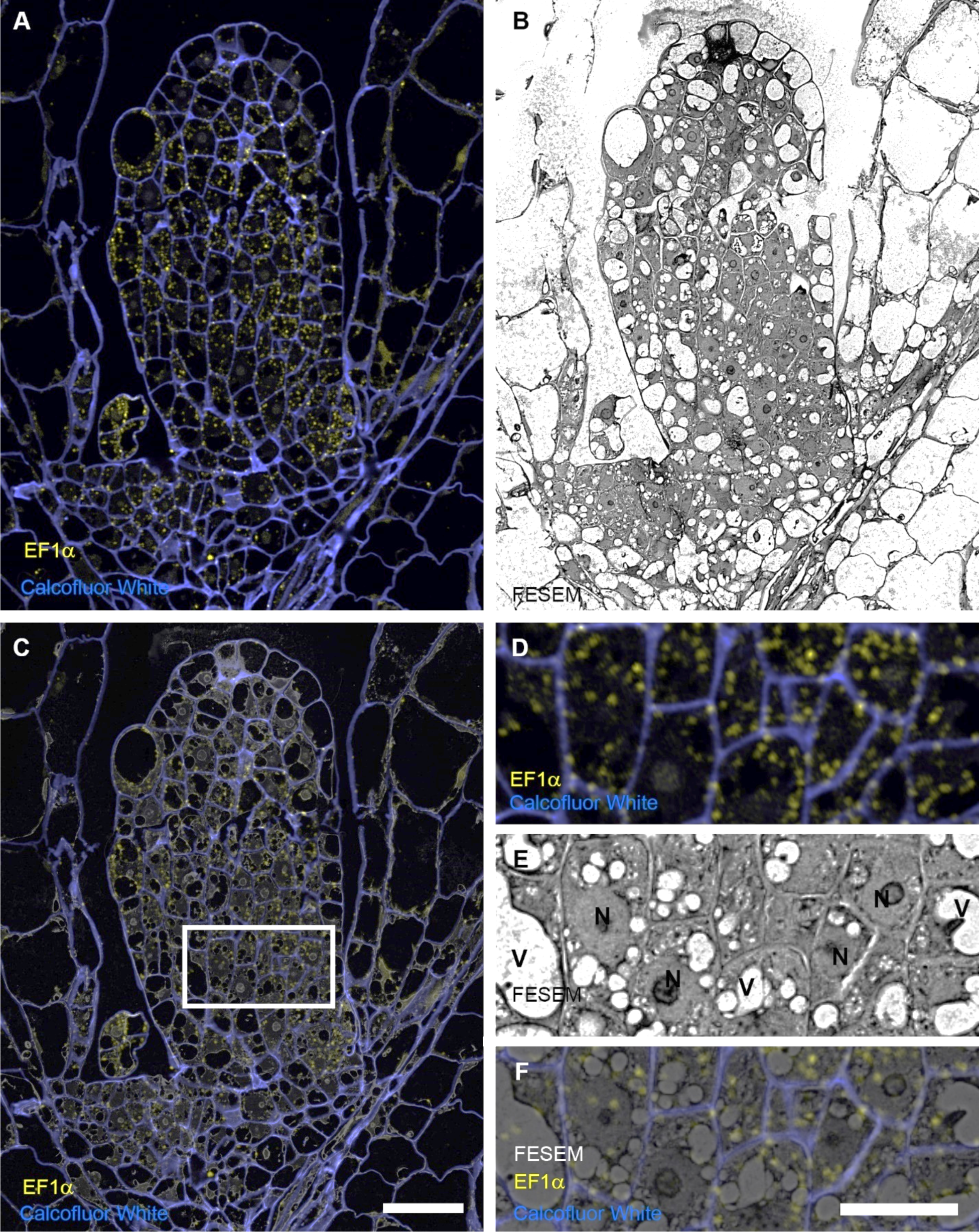
HCR can be combined with backscatter FESEM with methacrylate de-embedded plant tissues. (A) Fluorescence image of an *A thaliana* shoot meristem from 0.5 μM section stained with Calcofluor White (blue) and HCR EF1α (yellow) by confocal microscopy. (B) The same sample was heavy metal stained and imaged by FESEM using back-scatter electrons at 50nm pixel resolution for correlative microscopy. (C) The FESEM image intensities (from B) were then grayscale inverted and overlaid with the corresponding fluorescence mage (A) to yield a correlative light and electron microscopy image. The white box in C was enlarged to show corresponding fluorescence (D), FESEM (E) and combined correlated dataset (F) Scale in A to C = 50 µm, D to F = 20 µm.

## DISCUSSION

### Optimization of an HCR FISH protocol for plant sections

HCR FISH was initially developed for whole-mount animal tissues (15,23) and more recently also applied in whole-mount plant materials (25). Our study is the first to apply HCR FISH in plant sections, being suitable for a wide range of tissue types regardless of thickness and composition and opens the possibility to be combined with correlative ultrastructural studies. Here we provide protocols for both FFPE and methacrylate embedded tissues, as they have unique applications with inherent benefits and limitations. FFPE samples are routinely available and it’s straightforward to section large tissue areas and produce much thicker sections that generally contain a higher number of target mRNA molecules. However, FFPE samples generally provided poor sub-cellular morphological preservation and often had elevated levels of tissue autofluorescence. By contrast, methacrylate embedded, acetone de-embedded tissues demonstrated greatly improved morphological preservation but the reduced number of target molecules in semi-thin sections may limit their use to highly to moderately expressed genes without high quantum efficiency detectors. Additionally, sample sizes with methacrylate sections are relatively small (limited to a few millimeters) restricting the area of tissue that can be examined.

Cell wall digestion is required for probe penetration of whole-mount tissues, and proteinase K treatment is an essential step for revealing RNA targets in conventional ISH, which uses hapten-labeled long RNA probes (4,25,35). The split initiator HCR probes are un-modified 45 nt oligonucleotides and therefore can penetrate tissues more easily. Our experiments comparing treatments with and without proteinase K in both FFPE and methacrylate samples did not find a detectable difference in revealing HCR signal; thus, our protocol removed this step and the subsequent paraformaldehyde re-fixation required for conventional ISH, shortening the experimental time. More importantly, avoiding proteinase K treatment better preserved cellular morphology, allowing high-resolution detection of HCR signals at the cellular and subcellular levels and correlative study with FESEM.

### Autofluorescence and NSA in HCR FISH

Every HCR experiment should be run with tissue-only and hairpin-only negative controls to discern true signal from background autofluorescence and NSA as plants have numerous autofluorescent components. There are substantial differences in cell wall composition in grasses vs. dicots (grasses a lot more ferulates and coumarates than dicots), which affects fluorescence of the cell wall at some wavelengths. Furthermore, the plant cell wall contains molecules such as lignin, suberin and cutin, that are excited by UV or blue light, and emit violet to green light (36). We specifically avoided using the blue wavelengths for HCR and instead used it for detecting more intense cellulose staining using Calcofluor White or other UV stains, like DAPI for nuclei. Furthermore, Calcofluor White served as a useful alignment tool for correlative FESEM and potentially for multiplex studies. Ultimately, our protocol allowed for successful localization in our phylogenetically diverse test specimen.

Another abundant fluorescent molecule in plant tissues is chlorophyll, which is broadly excited, ranging from UV to red light, and emits red and far-red light (36). However, chlorophyll autofluorescence itself did not notably interfere with our experiments, probably due to chlorophyll removal by ethanol and acetone treatment during sample preparation and the fact that our probes were generally cytoplasmic. Generally, glutaraldehyde was avoided in the fixative as it enhances autofluorescence of proteins (36) and limits mRNA accessibility; however, very low levels of glutaraldehyde may be suitable for some labeling experiments, or where enhanced ultrastructural morphology is required (37).

Linear unmixing is a common method to remove background autofluorescence (38,39) and would certainly benefit plant studies. While we had limited access for use of linear unmixing in this study, we were able to create reference spectra of all tested fluorophores as well as characterize the spectral properties in *A. thaliana* seedling shoot meristems for linear unmixing. Our analysis showed strong green autofluorescence which decreased at lower energy wavelengths into the red and far-red spectrum. This knowledge is very useful when planning labeling experiments and matching anticipated signal levels from various probes relative to the autofluorescence background. Additionally, and consistent with this observation, punctate autofluorescence was observed in stem structures of the *Setaria* inflorescence at higher levels in Alexa Fluor 488 channel, moderate levels in the Alexa Fluor 546 channel but not detected in Alexa Fluor 647 channel (Fig. 2, Supp Fig. S1). Therefore, serving as a cautionary note, adequate controls described herein were mandatory to verify if non-specific puncta were an issue and if use of Alexa Fluor 647 was more appropriate.

Although the HCR probes are all 45 nt in length, probes produced by manufacturers are often non-pure and their lengths vary somewhat. If NSA controls show widespread signal in initial HCR experiments, the probes should be PAGE purified to obtain only full-length oligonucleotides around 45 nt. In our experience, using a glycerol-based antifade mounting medium, labeled sections should be imaged as soon as possible after completing the protocol but will usually maintain signal for about 2 – 3 weeks when stored at 4°C in the dark.

### Low intensity HCR FISH signal of ACT7

*ACTIN* genes are often used as housekeeping reference genes to normalize sample differences in differential gene expression quantification, for instance, by quantitative reverse transcription (1). ACT7 is highly expressed in rapidly expanding vegetative tissues, detected by promoter-GUS assays (40). Our inability to definitively detect ACT7 HCR signal with confocal microscopy may have been due to some probe pairs in the set failed to bind their target mRNA properly. Split-initiator probe design of HCR reduces non-specific binding to off-targets (23), but optimization of probes may be still required as in most ISH methods. Alternatively, transcripts in ACT7 may be more uniformly distributed in the plant cytoplasm, as opposed to local high-density clusters where intense foci were more conspicuously observed with *EF1*α**. Nevertheless, we did observe ACT7 HCR FISH signal above background controls using SIM super-resolution microscopy. This also demonstrated the value of high-sensitivity cameras and super-resolution microscopy for the detection of low expression levels and resulting fluorescence signals.

### Toward quantification in plant sections

While our efforts focused on developing the localization workflows that allowed for multiple probe classes to be combined with electron microscopy in plant tissues, a natural extension would be the potential for signal quantification. However, in our work, plant sections posed unique challenges in reliable quantification. First, consistency in section thickness must be considered and ensured to be uniform across samples. A housekeeping reference protein, gene or cell wall stain could be used for normalizing this variation but would need to be validated for the target tissues. Second, plant cells are highly variable in cell area and relative cytoplasm, often containing large central vacuoles (that take up ∼90% of the entire cell volume) and nearby air spaces that may bias the gross signal quantification per unit area. Thus, if quantification is intended, measurements should account for these morphological phenomena to avoid unintentional error. For example, a median section through a small dense cell in the meristem versus a large vacuolated mesophyll cell that may have the same labeling density in the cytoplasm itself, but the large central vacuole would suppress average signal intensity/total cell area. Third, as described herein, plant tissue and wavelength-specific autofluorescence may non-uniformly interfere with fluorescence signal quantification. Fortunately, when fixed emission filters and wavelengths were employed, far red wavelengths were universally more amenable to our tested cytoplasmic and cell wall probes. For greater flexibility, spectral strategies using linear unmixing from reference spectrum can be highly effective and were used successfully to alleviate the impact of autofluorescence in plant studies for quantitative FISH (41). While conventional ISH is semi-quantitative, some FISH-based techniques can be quantitative, including RNAscope® and HCR (24,42). Trivedi et al. (24) used two distinct sets of HCR probes labeled with spectrally separated fluorophores for the same mRNA targets and demonstrated that the two fluorescent signals are highly correlated in a linear relationship with an intercept of zero. Using a homozygous and heterozygous mouse embryo for a known gene, with the expectation that this gene will show 2:1 expression ratio, HCR showed approximately a two-fold difference in mRNA abundance, suggesting that HCR was quantitative (24). Further study and optimization will be required to determine if such quantitative approaches described in whole-mount or cell culture animal systems will translate to sectioned plant protocols.

## CONCLUSIONS

HCR FISH has emerged as a novel technique to map the spatial expression patterns of multiple genes concurrently. To apply this method in embedded plant tissues, we used *Arabidopsis* seedlings and *Setaria* inflorescence as our test systems with FFPE and methacrylate de-embedded plant sections. We found these preparations to be compatible with HCR and immunolabeling as well as correlative FESEM studies. In our approach working with sections, omitting proteinase K treatment did not notably affect signal detection and adequately preserved cellular structure. As expected, methacrylate sections produced much better cellular morphology at high resolution compared with paraffin, but working with sections that were often an order of magnitude thinner, required strategies to improve detection and sensitivity, especially with HCR. Despite some compromise in morphology with de-embedded methacrylate sections, adequate ultrastructure of the plant cells was possible after HCR and immunolabeling with thinner slices but was dependent on tissue and imaging conditions. As noted herein, numerous sources and patterns of plant autofluorescence can confound HCR and other affinity probe localization studies without careful assessment and adjustments to experimental design and/or technological strategies. However, several solutions, including judicious choice of fluorophore with labeled probes, leveraging available detection wavelengths, spectral unmixing strategies, instrument configuration and detector choice can help alleviate the problems. This work will serve as a useful basis for future improvements in labeling amplification, fixation, resin/resinless sectioning, multiplexing and potentially signal quantification. Our approach enables studying the localization of multiple RNAs, other affinity probes and the potential for cell pathways combined with cell ultrastructure to be assessed in plant systems as well as other organisms.

## MATERIALS AND METHODS

### Reagents and solutions

#### Reagents for tissue preparationC

Diethylpyrocarbonate (DEPC, Bioworld, Cat. No. 1609-47-8, Dublin, OH, USA) H_2_O: Add DEPC in milli-Q dH_2_O to a final concentration of 0.1%, stir overnight in a fume hood, and autoclave for 20 min.

10x phosphate buffered saline (PBS) in DEPC H_2_O: 1.37 M sodium chloride (NaCl), 27 mM Potassium Chloride (KCl), 43 mM Sodium Phosphate Dibasic (Na_2_HPO_4_), and 14mM Potassium Phosphate Monobasic (KH_2_PO_4_) in DEPC H_2_O, pH 7.3.

16% (w/v) paraformaldehyde (PFA) (Electron Microscopy Sciences, Cat. No. 15710, Hatfield, PA, USA).

0.2 M PHEM buffer (Electron Microscopy Sciences, Cat. No. 11165, Hatfield, PA, USA): 120 mM PIPES, 50 mM HEPES, 20 mM EGTA, 4 mM MgCl_2_, pH 7.2.

PFA fixative in PBS: 4% (w/v) PFA and 0.1% (v/v) Tween-20 in 0.1 M PBS. PFA fixative in PHEM: 4% (w/v) PFA in 0.1 M PHEM.

Histo-Clear® (National Diagnostics, Cat. No. HS-200, Atlanta, GA, USA).

Paraffin wax chips (Paraplast Plus, Cat. No. 39502004, McCormick Scientific, Saint Louis, MO, USA).

Embedding Ring (Simport Scientific. Cat. No. M460, Quebec, Canada) Base molds (Fisher Scientific. Cat. No. 22-363-553, Hampton, NH, USA)

Methyl Methacrylate/Butyl Methacrylate (Electron Microscopy Sciences, Cat. Nos. 18800 & 12100, Hatfield, PA, USA): Mix Methyl Methacrylate: Butyl Methacrylate in a 1:3 ratio. For final polymerization, under a stream of nitrogen gas, 0.5% (w/v) benzoin ethyl ether (Electron Microscopy Sciences, Cat. No. 11280, Hatfield, PA, USA) was added to the methacrylate monomer solution and mixed for at least 10 min, avoiding the introduction of oxygen in the resin, which can inhibit polymerization.

Vectabond® (Vector Laboratories, Cat. No. SP–1800, Newark, CA, USA)

#### Reagents for HCR

Ethanol series: 30%, 50%, 70% and 90% ethanol diluted in DEPC H_2_O.

Acetone (Sigma, Cat. No. 179124, St. Louis, MO, USA)

Nuclease free water (ThermoFisher, Cat. No. AM9930, Waltham, MA USA)

Deionized formamide (EMD Millipore, Cat. No. S4117, Burlington, MA, USA)

EDTA, 0.5M (VWR, Cat. No. 97062-654, Radnor, PA, USA))

0.2% glycine in PBS: Make 10% (w/v) Glycine in 1x PBS, dilute to a final concentration of 0.2% glycine with 1x PBS.

20x saline-sodium citrate (SSC) in DEPC H_2_O: 3 M NaCl, 0.3 M sodium citrate in DEPC H_2_O.

Probe Hybridization Buffer (Molecular Instruments, Los Angeles, CA, USA): 30% formamide, 5x SSC, 9 mM citric acid (pH 6), 0.1% Tween-20, 50 µg/ml heparin, 1X Denhardt’s solution and 10% dextran sulfate.

50% Probe Wash Buffer: Dilute Probe Wash Buffer (Molecular Instruments, Los Angeles, CA, USA) in 5x SSCT (0.1%Tween-20). The final concentrations are 15% formamide, 750mM sodium chloride, 75mM sodium citrate, 4.5 mM citric acid (pH 6), 0.1% Tween-20 and 25 µg/ml heparin.

Amplification Buffer (Molecular Instruments, Los Angeles, CA, USA): 5x SSCT (0.1%Tween-20) and 10% dextran sulfate.

Hairpins (H1 and H2) with conjugated fluorophores (Alexa Fluor 488/546/647) for each system used (B1 – B5, Molecular Instruments, Los Angeles, CA, USA)

Oligonucleotide Probe Sets (IDT, Coralville, IA, USA or Molecular Instruments)

#### Solution used for probe PAGE purification and in vitro assays

Formamide loading buffer: deionized formamide (Sigma-Aldrich, Cat No. F9037, Burlington, MA, USA) with 10 mM EDTA

Elution buffer: 500 mM ammonium acetate, 10 mM magnesium acetate, 1 mM EDTA, 0.1% SDS, pH 8.0

1x TBE buffer: 89 mM Tris base, 89mM boric acid, 2mM EDTA

GelRed (Phenix Research Products. No.C755G19, Swedesboro, NJ, USA) RedSafe (Bulldog Bio, Woburn, MA, USA)

100 bp Plus and 1 kb Plus DNA ladders (Goldbio, Saint Louis, MO, USA)

6X loading buffer (New England Biolabs, Ipswich, MA, USA)

#### Reagents for Immunolabelling

Rabbit anti-actin polyclonal antibody (Agrisera, Cat. No. AS 13 2640, Vannas, Sweden)

LM19 rat anti-homogalacturonan monoclonal antibody (Agrisera, Cat. No. AS18 4191, Vannas, Sweden)

Donkey anti-rabbit IgG DyLight 650 (Agrisera, Cat. No. AS12 2329, Vannas, Sweden) Goat anti-rat IgG Alexa Fluor 647 (Invitrogen, Cat. No. A21247, Vannas, Sweden)

Blocking buffer: 5% bovine serum albumin (BSA) (Sigma-Aldrich, Cat. No. A8577, Saint Louis, MO, USA) in PBST (0.1% Tween-20)

#### Reagents for fluorescent imaging

Calcofluor White solution: 5 mM Calcofluor White (Biotium, Cat. No. 29067, Fremont, CA, USA) to a final concentration of 50 µM in water.

DAPI (4′,6-diamidino-2-phenylindole) solution: Dilute 1 mg/ml DAPI (ThermoFisher, Cat. No. 62248, Rockford, IL, USA) to a final concentration of 0.5 µg/ml in 1x PBST (0.1% Tween-20).

SlowFade^TM^ Glass antifade mountant, 1.52 refractive index (Thermo Fisher Invitrogen, Cat. No. S36917, Waltham, MA, USA)

Two-component silicone-glue Twinsil® (Picodent, Reagent A Cat. No. 32002.42011-12, Reagent B Cat. No. 34001.02014-06 Wipperfurth, Germany): Mix the two components in 1:1 ratio.

#### Reagents for FESEM

2% Osmium tetroxide (Electron Microscopy Sciences, Cat. No. 19110, Hatfield, PA, USA): dissolve two 1-g ampules in 100 ml of water.

0.1% KMnO_4_ in 0.1 N H_2_SO_4_: Add 0.01g KMnO_4_ (Sigma-Aldrich, Cat. No. 223468, St. Louis, MO, USA) in 10 mL 0.1 N H_2_SO_4_ (Fisher Scientific, Cat. No. SA818-500, Pittsburg, PA, USA). This solution was filtered with a 0.2 µm PES filter (Whatman UnifloTM, Cytiva, Buckinghamshire, UK) prior to use.

1% (w/v) methanolic uranyl acetate: Add 1g uranyl acetate (SPI Supplies, Cat. No. 02624-AB, West Chester, PA, USA) to 100 mL methanol (Electron Microscopy Sciences, Cat. No. 18510, Hatfield, PA, USA): This solution was filtered with a 0.2 µm PES filter prior to use.

Sato’s lead solution: Add 0.20 g anhydrous lead citrate (Electron Microscopy Sciences, Cat. No. 17800, Hatfield, PA, USA), 0.15 g lead nitrate (Electron Microscopy Sciences, Cat. No. 17900, Hatfield, PA, USA), 0.15 g lead acetate (Electron Microscopy Sciences, Cat. No. 17600, Hatfield, PA, USA), 1.00 g sodium citrate (Electron Microscopy Sciences, Cat. No. 21140, Hatfield, PA, USA) and 41.0 mL distilled water in a 50 mL volumetric flask, mix well and adjust pH with 9 mL 1N NaOH (Electron Microscopy Sciences, Cat. No. 21160, Hatfield, PA, USA). This solution was filtered with a 0.2 µm PES filter prior to use.

### Equipment and Supplies

- Leica EG1150 tissue embedder
- Leica RM2255 rotary microtome
- Incubators set to 37°C and 60°C
- Slide warmer
- Probe-On Plus Slides (Fisher Scientific, Pittsburgh, PA, USA)
- Adhesive treated coverslips: Glass coverslips (No. 1.5, 22mm x 22mm) were coated with Vectabond® (Vector Laboratories, Newark, CA, USA) following the manufacturer’s recommendations and allowed to dry.
- EMS UVC3 Ultraviolet Cryo Chamber (Electron Microscopy Sciences, Cat. No. 70445-10, Hatfield, PA, USA)
- BEEM capsules (Size 00, 8mm I.D, Electron Microscopy Sciences, Cat. No. 70000-B)
- DiATOME Ultra 45°, 2.0 mm Diamond knife (Diatome, Quakertown, PA, USA
- Leica UltraCut UCT ultramicrotome (Leica Microsystems, Vienna, Austria)
- Azure Biosystems C6 Gel Scanner (Dublin, CA, USA)
- Heat block set to 95°C
- Vacuum desiccator
- Sealed humid container (This can be made in-house from many types of plastic containers which have a sealable lid. Elevating the glass slides and coverslips off the bottom with spacers helps to avoid the solutions wicking away from the sections into the moist paper towels below)
- Microfuge
- NanoDrop spectrophotometer
- Glass Coplin jars
- Slide mailers
- Coverslip mini-rack (Thermo Fisher, Cat. No. C14784, Waltham, MA, USA)
- 50 mLbeakers
- Parafilm (Bemis, IL, USA)
- Leica TCS SP8 confocal laser scanning microscope (Leica Microsystems, Wetzler, Germany)
- ZEISS Elyra 7 super-resolution microscope with Lattice SIM² (Zeiss, Jena, Germany)
- ZEISS Merlin field emission scanning electron microscope with ATLAS 5 Software (Zeiss, Oberkochen, Germany)
- ZEISS LSM980 confocal microscope with Quasar spectral detector (Zeiss, Jena, Germany)

### Plant Materials

*Setaria viridis* (accession A10) seeds were sown in Jolly Gardener soil, kept at 4°C in the dark for 2 weeks to break dormancy, then grown in a growth chamber maintaining 12 h/12 h (day/night) photoperiod and 31°C/22°C (day/night) temperatures, 50% humidity and 350 µmol/m^2^/sec light intensity. *Arabidopsis thaliana* (Col-0) seeds were sterilized and sown on ½ strength Murashige & Skoog medium, 1% sucrose and 0.8% agar (pH 5.8) in a growth chamber with a 16 h/8 h day/night (22°C/19°C) regime, 50% humidity and 200 µmol/m^2^/sec light intensity.

### HCR and Immunofluorescence Sample Processing and Imaging Workflow

We encapsulated the most complete scenario for labeling and imaging workflows of multiple probe classes in Figure 8. The vast majority of the processing time involves sample fixation and embedment and generating tissue sections. Immunolabelling and correlative FESEM can follow HCR and are optional. Probe sets that produce relatively low signals (below the detection limit of laser scanning confocal microscopy (LSCM)) can be successfully imaged with high quantum efficiency imaging modalities. The details of the experimental workflow are as follows below:

**Figure 8.**
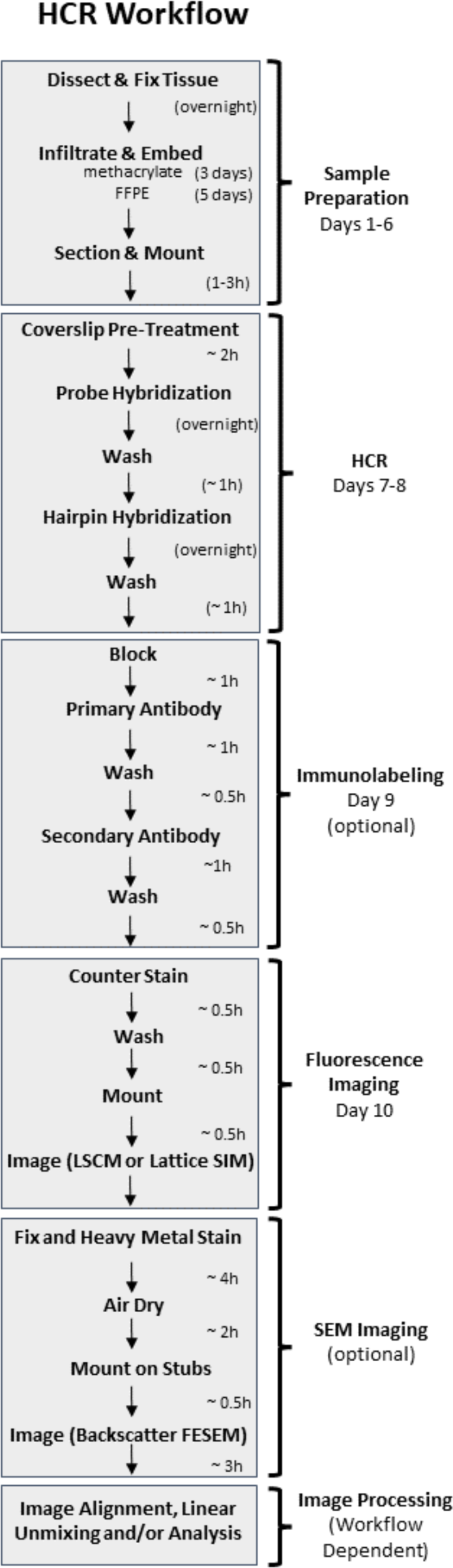
Representative HCR workflow and durations. The majority of experimental time involves preparation for sample fixation and generating tissue sections. Immunolabelling following HCR and/or SEM for correlative ultrastructure is optional. Probe sets that produce relatively low signals (below the detection limit of laser scanning confocal microscopy (LSCM)) may be imaged with super-resolution lattice structured illumination microscopy (SIM) or other high quantum efficiency imaging modalities.

### HCR Probe Set Design

Custom split initiator probe sets may be designed and ordered directly from Molecular Instruments or manually designed based on the specifications described in (23) as follows. A list of prospective 22 nt oligomer probes was generated using an online probe design tool (43), and their locations were manually mapped onto the target sequence using the free plasmid DNA mapping software SnapGene Viewer. Regions of the target mRNA where two 22 nt probes were located in close proximity (20 nt or less) were considered as prime locations for generating HCR probe pairs. The sequences of the probes were then manually extended to 25 nt each, and their locations were adjusted so that the pairs were separated by 2 nt of either G and/or C and had predicted melting temperatures of 35-60°C for each probe. This process was repeated until probe pairs were spaced along the entire CDS and 3’UTR of the target mRNA. The probe sequences were confirmed to be specific using NCBI nucleotide BLAST. Final probe sets were composed of 6-12 probe pairs (12-24 probes). One of the five unique initiator sequences (B1-B5) (23) was chosen for each target mRNA probe set. The 5’ or 3’ half of the 36 nt initiator sequence followed by a 2 nt spacer was added to either the 5’ end of the odd numbered anti-sense probe sequence, or the 3’ end of the even numbered anti-sense probe sequence, respectively, resulting in a final HCR probe length of 45 nt. The commercially generated oligomers with standard desalting were dissolved to 100 µM in nuclease free water. The individual stock probe solutions were then mixed and diluted to 2 µM/probe in nuclease free water. Initiators and full sequences of the probe sets used in this work are shown in Table S1.

### HCR Probe PAGE Purification (1 day)

Probe sets which showed non-specific puncta on an initial HCR experiment were PAGE purified as follows. An 8% polyacrylamide (8 M urea, 1x TBE) solution was mixed, degassed for 15 min under vacuum and polymerized with 0.05% fresh ammonium persulfate and 0.001% TEMED and poured into two 1mm Bio-Rad Protean II mini gels. The gels were pre-run at 200V for 30 min in 1x TBE buffer. 100 µg total of the odd numbered probe stocks were mixed and diluted with an equal volume of formamide loading buffer, heated to 95°C for 2 min, then snap cooled on ice. The entire volume of the odd numbered probe mixture was loaded equally into the wells of one gel along with a single well of formamide loading buffer containing 0.1% xylene cyanole FF and 0.1% bromophenol blue to serve as size markers. The even numbered probes were prepared in a similar fashion and loaded into the second gel and both were run at 200V until the bromophenol blue reached the bottom of the gel. The edges of the gels were notched at the locations of the two dye bands (xylene cyanole 75 bp, bromophenol blue 19bp), stained with GelRed (Phenix Research Products), imaged under UV and the large 45 nt band isolated, minced and loaded into 0.5 ml microfuge tubes. The tubes were perforated by a sharp needle five times, inserted into 1.5 mL microfuge tubes, and spun at 10,000 g until the gel had been crushed into the larger tubes. Next, 400 µl of elution buffer was added to the gel pieces and incubated at 37°C overnight. The elution buffer with DNA was removed from the gel pieces, and the DNA precipitated with 1/10th volume of 3 M sodium acetate and 2.5 volumes of 100% ethanol at −20℃ for 30 min. The DNA was pelleted at 10,000 x g for 30 min at 4℃, and the supernatant was removed. The pellet was washed in 70% ethanol at RT, dried at RT for 15 min and dissolved in nuclease free water. Probe concentrations were determined by a NanoDrop spectrophotometer.

### In-vitro HCR Assay (optional)

A custom 52nt oligo, corresponding to the sense strand of the CAB1 mRNA targeted by probe 1 and 2, was generated commercially (IDT, Coralville, IA) along with the 36 nt system B2 initiator sequence (Table S1) (23). The B2 hairpins, conjugated to Alexa Fluor 546, were heated to 95°C for 90 sec then allowed to cool at RT in the dark for 30 min. The target, initiator and probes were heated to 95°C for 90 sec then snap cooled on ice. Next, 12 µl reactions were then set up in 5X SSCT (0.1% Tween-20) with the following final concentrations: System B2 hairpin H1 and H2 each 125 nM, B2 initiator 5 nM, target oligonucleotide 10 nM, split initiator probes 1 and 2 at 10 nM each. The reactions were incubated at 25°C for 4 hr. 2µl of 6X loading buffer (New England Biolabs, Ipswich, MA) was added to each reaction and then loaded onto a 1% agarose/TBE gel along with 5 µl each of 100 bp Plus and 1 kb Plus DNA ladders (Goldbio, Saint Louis, MO). The gel was run at 105V for 45 min and stained with 1X RedSafe (Bulldog Bio, Woburn, MA) in TBE for 30 min. The gel was imaged with an Azure Biosystems C6 Gel Scanner (Dublin, CA) with peak excitation/emission wavelengths of 530/572 nm.

### Fixation, Embedding and sectioning (5-6 Days)

Once HCR Probes are generated and purified, all subsequent steps below provide details from specimen fixation until image processing as outlined for the HCR Workflow (Fig.8).

After 17 days in the growth chamber, *Setaria* panicles were dissected under a stereo microscope. Panicles were then fixed in PFA fixative in PBS under vacuum for 10 min on ice, then overnight in fresh fixative at 4°C. All processing steps were conducted in standard 20 mL glass scintillation vials. After washing two times in 1x PBS for 30 min each on ice, the tissue was processed through a standard paraffin embedding protocol. The samples were dehydrated through a graded series of ethanol (30%, 50%, 70%, 95%, 100%, 100%) for 1 hr each on ice, followed by 100% ethanol overnight at 4°C. The ethanol was gradually replaced using 1 hr incubations in 25%, 50%, 75% Histo-Clear® diluted in ethanol at 22°C (RT), followed by three times in 100% Histo-Clear^®^. Then the vial was filled with paraffin wax chips and incubated overnight at RT, followed by 2 h incubation at 42°C and at 60°C until the wax chips completely melted. The solution was replaced with fresh 100% melted paraffin at 60°C six times over 2-3 days. The samples were then embedded in paraffin using a Leica EG1150 tissue embedder, allowed to harden and stored at 4°C. Next, 10µm longitudinal sections were cut using a Leica RM2255 rotary microtome, placed on Probe-On Plus Slides, briefly stretched on a droplet of water and allowed to dry overnight on a 37°C slide warmer. Slides were stored desiccated at 4°C in the dark.

Three-day-old *Arabidopsis* seedlings were fixed in PFA fixative in PHEM buffer at 4°C. Following the ethanol dehydration series, half the seedlings were processed in paraffin (see above), while the remaining samples were embedded in methacrylate resin. The seedlings were infiltrated in 25%, 50%, 75% methacrylate (1:3 methyl methacrylate: n-butyl methacrylate) diluted in ethanol for 2 h each, followed by two changes of 100% methacrylate for 24 h each at RT. The seedlings were then embedded in methacrylate with benzoin ethyl ether under a constant stream of nitrogen gas in BEEM capsules and polymerized under UV radiation for 48 h at −20°C using a UVC Cryo Chamber filled with dry ice. Capsules were removed and the blocks allowed to cure in a fume hood at RT until solid. 0.5-1.5 µm semi-thin sections were cut using a diamond knife and Leica UltraCut UCT ultramicrotome. The sections were stretched with chloroform vapors and transferred to adhesive-treated coverslips to allow drying at 42°C for 1h. Coverslips were stored at 4°C in the dark. The best experimental results were obtained using sections cut within ∼2 weeks of the HCR procedure.

### HCR (2.5 days)

Paraffin sections on slides were incubated for 2 x 10 min in Histo-Clear^®^ in glass Coplin jars to remove the wax. Methacrylate sections on coverslips were de-embedded for 10 min in acetone using a coverslip mini-rack in a 50 mL glass beaker. For all remaining steps, the paraffin and methacrylate sections were treated in parallel. Wash steps for the paraffin sections were conducted in plastic slide mailers. Methacrylate sections continued to be processed by moving the coverslip mini-rack through a series of glass beakers with the appropriate solutions.

Sections were rehydrated in a decreasing graded series of ethanol (100, 90, 70, 50 and 30%) for 2 min each, followed by two washes in 1x PBS for 2 min each. Sections were then incubated in 0.2% glycine in PBS for 5 min, washed in 1x PBS twice for 2 min each and dehydrated in a series of increasing graded ethanol for 2 min each, ending in 2 changes of 100% ethanol for 5 min each. Sections were air dried in a vacuum for 1 h at RT. This pre-treatment regime eliminates several steps commonly found in conventional *in situ* hybridization protocols including proteinase K digestion, re-fixation and triethanolamine (TEA) incubation. These steps had no notable effect on HCR labeling results and eliminating them resulted in a shorter and simpler protocol as well as reduced extraction that would impact morphology and some antigens for immunolabeling.

Custom-designed split initiator oligonucleotide probes were diluted in the hybridization buffer at a final concentration of 20 nM/probe. The probe solution (125 µL for slides and 75 µL for coverslips) was pipetted over the sections and covered with a piece of Parafilm which had been cut to a slightly smaller dimension than the slides or coverslips. The slides and coverslips were incubated at 37°C overnight in a moist chamber. The hybridization solution was drained away, and the slides and coverslips were washed in 50% Probe Wash Buffer at 37°C for 15 min, followed by 2x SSCT (0.1% Tween-20) incubation at 37°C for 15 min and a final 15 min wash in 5x SSCT (0.1% Tween-20) at RT.

Each pair of fluorophore-conjugated hairpins (H1 and H2) needed for an experiment was pipetted into separate microfuge tubes and heated to 95°C for 90 sec in a heat block, then allowed to refold at RT for 30 min in the dark. The tubes were then spun down and the hairpins were added to Amplification Buffer at a final concentration of 60 nM. The hairpin buffer was then pipetted over the sections, covered with Parafilm, and incubated in a moist chamber overnight at RT in the dark. The hairpin buffer was drained off, and the slides and coverslips were washed 3 x 10 min in 5x SCCT.

### Immunolabeling (0.5 day)

In experiments where immunolabeling was performed after HCR, the sections were incubated in blocking buffer for 1 h at RT in a moist chamber. Primary antibodies (125 µL per slide and 75 µL per coverslip) were diluted in blocking buffer (1:15 dilution for LM19 and 1:200 dilution for anti-actin) and incubated on the sections for 1 h in the dark at RT in a moist chamber. The primary antibody solution was drained off and the sections were washed 3 x 15 min in blocking buffer.

Fluorophore-conjugated secondary antibodies were diluted 1:1000 in blocking buffer and incubated on the sections for 1 h at RT in the dark in a moist chamber. The secondary antibody solution was drained off and the sections and coverslips were washed 3 x 15 min in PBST.

### Cell Wall and Nuclear Staining (∼0.5 h)

After HCR (and immunolabeling), samples were stained to differentiate individual cells. Cell walls were stained with Calcofluor White diluted 1:200 in water for 20 min at RT in the dark in a moist chamber. Nuclei were stained with DAPI solution for 30 min at RT in the dark in a moist chamber. The slides and coverslips were washed in 1x PBS, quickly dipped into water, mounted using SlowFade^TM^ Glass antifade mountant, and sealed with Twinsil. The samples can be imaged immediately or stored at 4°C in the dark for up to 2 weeks without appreciable loss of HCR signal or specificity.

### Imaging and Image Processing

All the laser scanning confocal microscope images were taken with a Leica TCS SP8 with white light lasers using the Leica Las X software and the following objectives: 63x/1.2 HC PL APO CS2 water immersion, 40x/1.1 PL APO CS2 water immersion and 20x/0.7 HC PL APO air lens. Images were taken at 1024 x 1024 pixels with a bit depth of 12. Calcofluor White was excited using the 405 nm diode laser line and collected at 410-473 nm. Alexa Fluor 488 was excited using the 488 nm laser line and collected at 508-552 nm. Alexa Fluor 546 was excited using the 514 nm laser line and collected at 571-628 nm. Alexa Fluor 647 and DyLight 650 were excited with the 664 nm laser line and collected at 662-748 nm. Spectral imaging was performed using a Zeiss C-Apochromat 40X/1.2 W Korr FCS objective on a ZEISS LSM980 equipped with a Quasar spectral detector using simultaneous laser excitation with 405 nm, 488 nm, 561 nm and 640 nm diode laser lines and covering 411-694 nm with 32 channels at 9 nm GaAsP-PMT detector windows. Images were acquired at 16 Bit, 69 nm x-y pixel resolution, 3091 x 3091 pixel frame size and brightness levels were adjusted uniformly for visualization using Fiji (44) and the images were converted to TIFF format for final figure assembly. Experimental and negative control images were taken using identical microscope settings and displayed with matching black and white levels with experimental samples in Fiji. Lattice SIM super-resolution images were taken with a Zeiss Elyra 7 inverted microscope running the Zen Black 3.0 software package and a 40x/1.4 Plan Apochromat DIC M27 oil immersion objective. Consecutive z-stack images were taken at 1024×1024 pixel frame size and at 16 Bit. Voxel scaling was 0.0099 x 0.099 x 0.329 μm. Calcofluor White was excited with 405-nm and ACT7 with 488-nm lasers sequentially in fast frame mode. Respective emissions were detected using the CMOS camera and BP420-480 + BP 495-550 filter. The Zeiss SIM^2^ deconvolution processing module was used to process the images and option “scale to raw image” was selected to retain original relative signal intensities.

### Staining coverslips for SEM (∼3 h)

This protocol was modified from (45). Coverslips were placed section-side up on a petri dish and rinsed well with double-distilled water (ddH_2_O) on a benchtop shaker at low speed for 15 mins. Then in a fume hood, the coverslips were floated section-side down on 200µL droplets of 2% Osmium tetroxide for 1 hour and rinsed with ddH_2_O for 10 mins. Coverslips were then floated on droplets of 0.1% KMnO_4_ dissolved in 0.1N H_2_SO_4_ for 1 min and then rinsed with ddH_2_O for 5-10 mins. Drops of filtered 1% methanolic Uranyl Acetate (UA) were placed in a petri dish lined with parafilm and the coverslips were floated on the UA droplets for 30 mins. Next, the coverslips were rinsed by floating on drops of methanol for 5 mins and were allowed to dry before staining with lead. The lead staining chamber was prepared by adding NaOH pellets in a petri dish and the coverslips were floated on drops of filtered Sato’s lead solution (46) for 1 min. Finally, coverslips were then rinsed using degassed ddH_2_O and excess water was carefully blotted dry and coverslips were allowed to air dry.

### SEM imaging

The dried coverslips with sections facing up were mounted onto 25mm aluminum pin stubs with double-sided adhesive carbon tabs and then conductive silver paint was applied connecting the aluminum stub to the top edge of the coverslip surface, ensuring a conductive trail. After the conductive paint dried, the entire coverslip was carbon coated prior to imaging. A ZEISS Merlin was used at 3kV and scanned at 50 nm or 5 nm pixel resolutions and 8 Bit depth with back-scatter electrons detector and with FIBICS ATLAS 5 (v5.3.5.27). Confocal and FESEM images were overlaid using Adobe Photoshop CC 2019, Linear Dodge (Add) was applied to the confocal image to see the combined back-scatter EM and fluorescence image, The Free Transform tool was used to rotate, rescale and align the fluorescence image with the corresponding FESEM image using the plant cell wall (Calcofluor White from the confocal image) and FESEM images for registration features. The Warp non-linear transformation function was used for the final, very minor local cell wall changes that occurred between confocal and FESEM sample treatments.

## List of abbreviations

Abscission zone: AZ.
*ACTIN7*: *ACT7*.
*CHLOROPHYLL BINDING FACTOR a/b*: *CAB1*.
Digoxygenin: DIG.
*ELONGATION FACTOR 1 ALPHA*: EF1*α*.
FFPE: Formalin fixed Paraffin Embedded.
Field emission scanning electron microscopy: FESEM.
FISH: fluorescent *in situ* hybridization.
HCR: hybridization chain reaction.
KNOTTED 1: KN1.
Nitroblue tetrazolium/5-bromo-4-chloro-3-indolyl-phosphate: NBT/BCIP.
PAGE: polyacrylamide gel electrophoresis.
Rolling circle amplification: RCA.
SHATTERING 1: SH1.
Single molecule FISH: smFISH.
SIM: Structured Illumination Microscopy.
Transcription factor: TF.
Tyramide signal amplification: TSA.
Uranyl acetate: UA.

## Declarations

### Availability of data and materials

The transcript sequences of *Setaria viridis SH1* (Sevir.5G293800.1), *KN1* (Sevir.9G107600.1) and MYB26 (5G293800.1) are available on Phytozome 13 (https://phytozome-next.jgi.doe.gov/). The transcript sequences of *Arabidopsis thaliana CAB1* (AT1G29930), *EF1*α** (AT1G07940) and *ACT7* (AT5G09810) are available on TAIR (https://www.arabidopsis.org). The probe sequences and images are available within the paper and supplemental materials.

### Authors’ contributions

KC, YY, SB, ML, EAK and BM obtained funding. KC, YY, SB, ML designed the research. KC, DH and YY designed the experimental approach. SB, DH and YY prepared the plant materials. DH performed the HCR and Immunolabeling experiments. YY performed the colorimetric ISH. DH, KC and AK acquired the images. JW and KC performed the FESEM. DH, ML and KC performed the image analysis. DH, YY, EAK and KC drafted the manuscript, and all authors discussed the results and edited the manuscript.

## Acknowledgements

We gratefully thank the NSF for funding this work (Award # 2130365 to K. Czymmek) and Joanna Porankiewicz-Asplund and Agrisera for providing our tested DyLight Secondary antibodies. We also acknowledge imaging support from the Advanced Bioimaging Laboratory (RRID:SCR_018951) at the Danforth Plant Science Center and usage of the ZEISS Elyra 7 Super-Resolution Microscope acquired through an NSF Major Research Instrumentation grant (DBI-2018962). We are also especially appreciative of the expert ZEISS FESEM and ATLAS support from Dr. Sanja Sviben and Washington University Center for Cellular Imaging (WUCCI).

---

**Figure S1.**
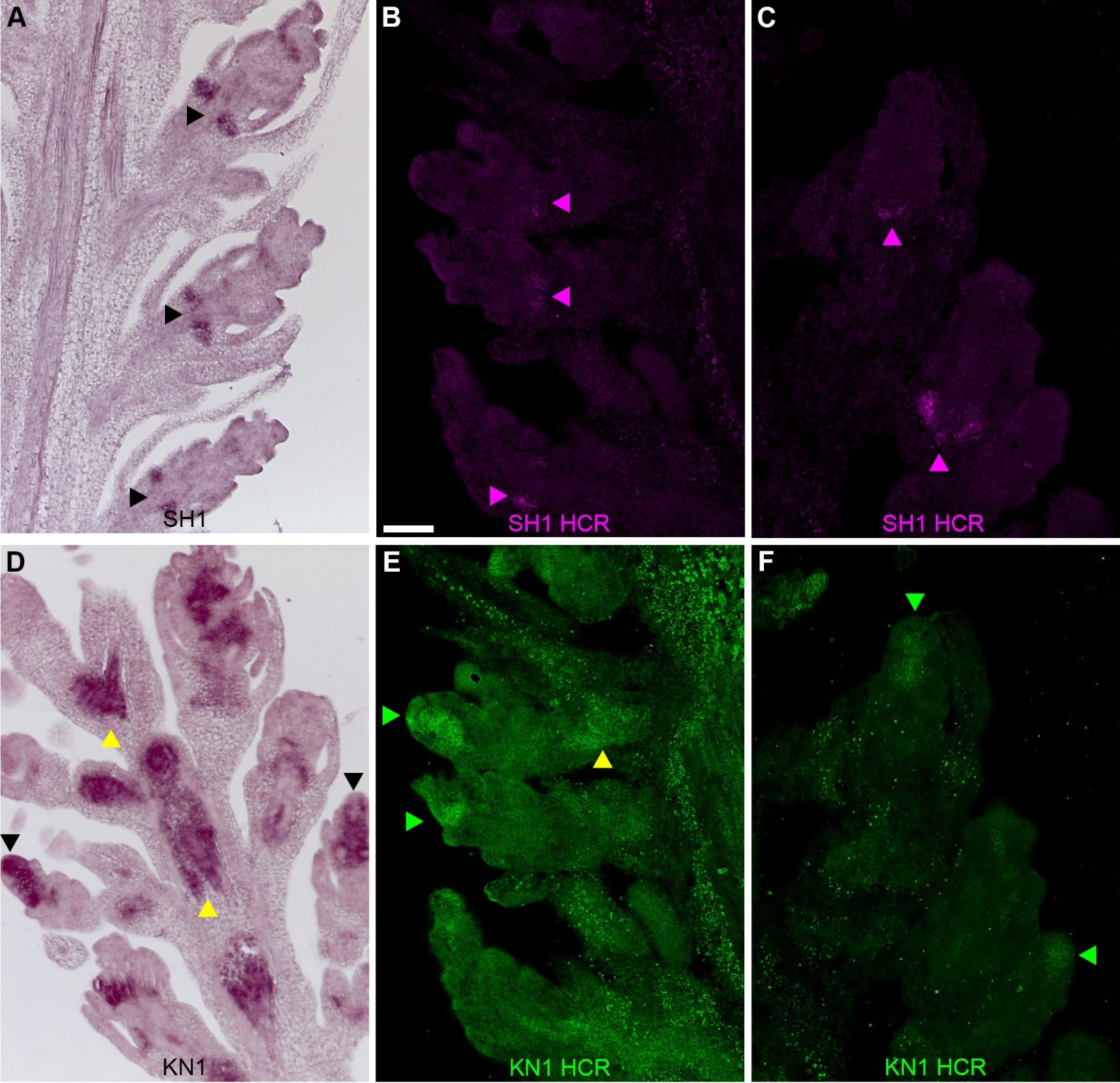
Localization patterns for SH1 and KN1 in parµμµfin embedded *Setaria viridis* spikelets are consistent using both HCR and conventional ISH. A-F. 10 µm paraffin longitudinal sections through the panicle of *Setaria viridis*. A. Light micrograph of colorimetric ISH against Shattering 1 (SH1). Black arrows indicate SH1 expression in the abscission region of three spikelets. B-C. Confocal images of HCR detection of SH1 using Alexa Fluor 546 conjugated B2 hairpins. Magenta arrows indicate SH1 transcript expression in the abscission zone of multiple spikelets. D. Light micrograph of colorimetric ISH against the gene *Knotted* 1 (KN1). Black arrows indicate KN1 expression in the meristem region of two spikelets. KN1 expression is also seen in the vascular meristem (yellow arrows). E-F. Confocal microscope images of HCR detection for KN1 using Alexa Fluor 488 conjugated B4 hairpins. Green arrows indicate KN1 transcript expression in the rounded meristem tissue of multiple spikelets. Yellow arrows indicate KN1 expression in the vascular tissue. A pixel intensity threshold was applied to reduce the saturated non-specific puncta in E-F. Scale = 50 um.

**Figure S2.**
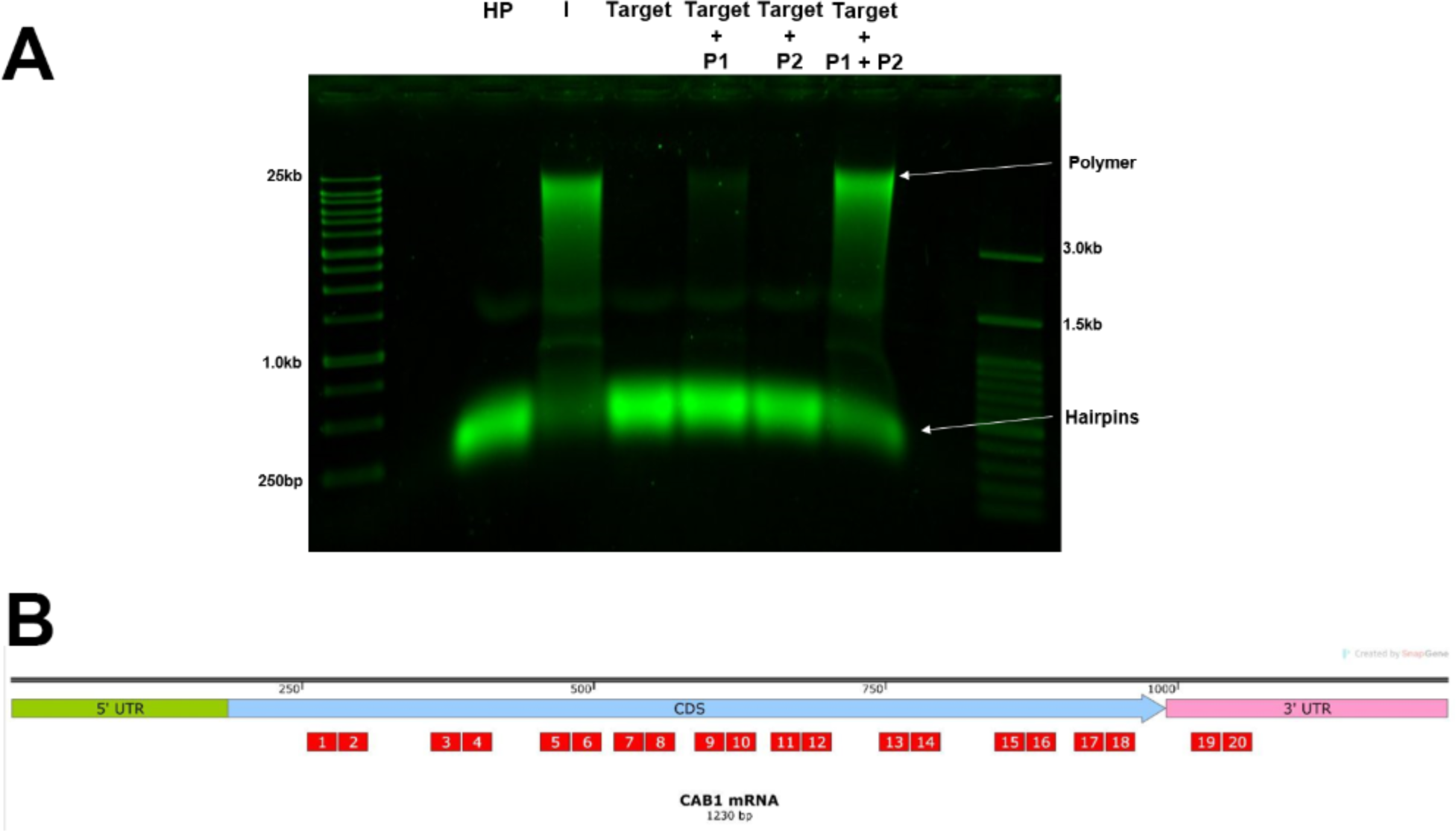
*In vitro* hairpin assembly. A. *In vitro* assay confirming the formation of hairpin polymers only when both split initiator probes in a pair hybridize to the CAB1 target mRNA. HP: hairpins only. I: B2 initiator sequence. P1: CAB1 probe 1. P2: CAB1 probe 2. 1% agarose/TBE gel with DNA ladders. B. Map of the mRNA sequence of *Arabidopsis* CAB1. Red boxes indicate the positions of the HCR probe pairs along the target transcript.

**Fig. S3.**
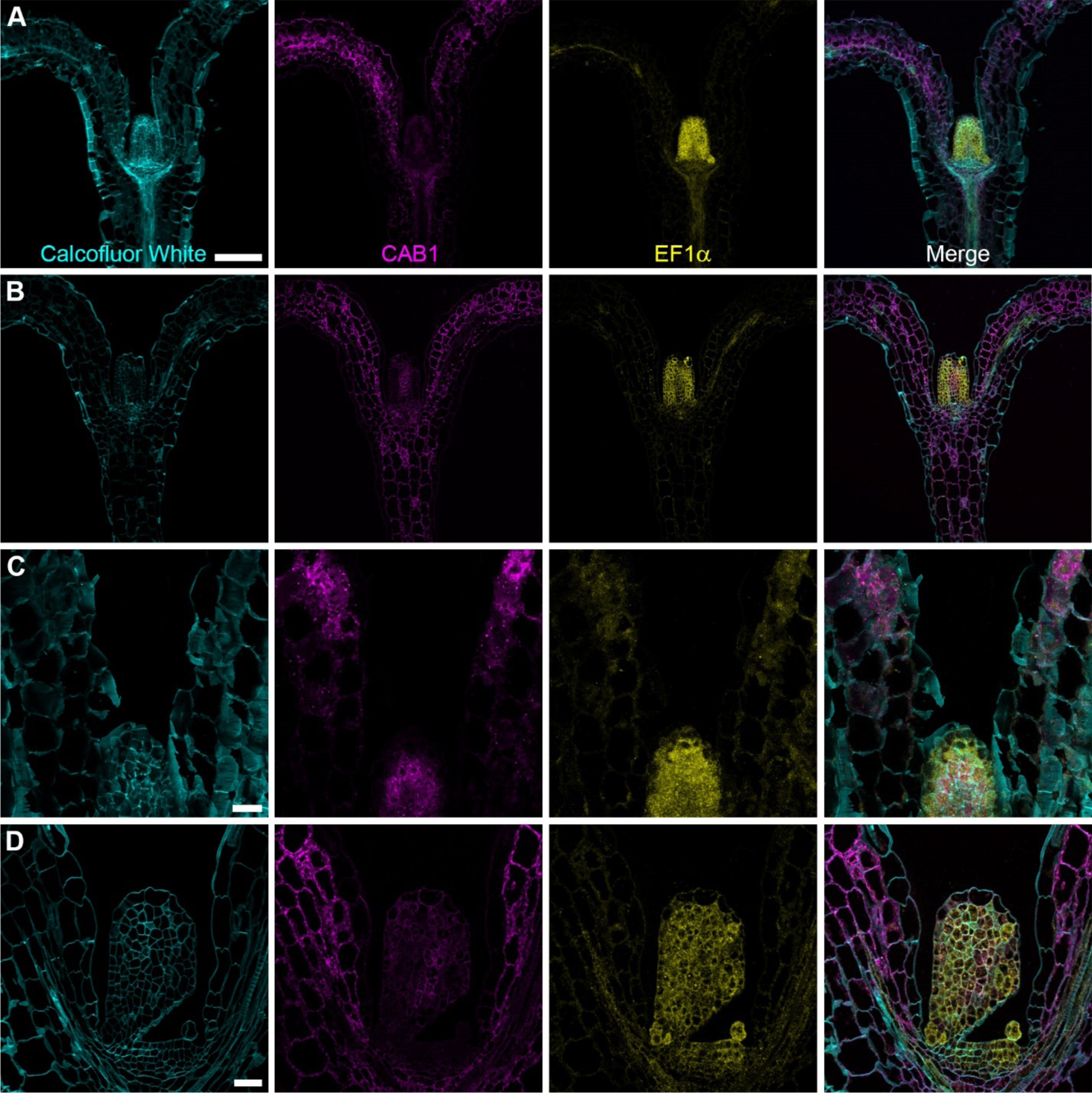
Comparison of HCR in paraffin and methacrylate sections. A-B. 3-day *Arabidopsis thaliana* seedling cotyledons, hypocotyl and shoot apex sectioned in the longitudinal plane and imaged using a 20x/0.7NA objective. MIP of Z stack. A. 10 µm paraffin section. Scale = 100 µm. B. 1.5 µm methacrylate section. C-D. 3-day *Arabidopsis thaliana* seedling cotyledons and shoot apex sectioned in the longitudinal plane and imaged using a 63x/1.2NA water immersion objective. Single 1.37 µm optical slice. C. 10 µm paraffin section. D. 1.5µm methacrylate section. Scale = 20 µm. HCR for CAB1 (Alexa Fluor 546) and EF1a (Alexa Fluor 647) with calcofluor white as a cell wall marker. CAB1: Chlorophyll binding protein a/b. EF1*α*: Elongation Factor 1 alpha. Scale = 20 µm.

**Figure S4.**
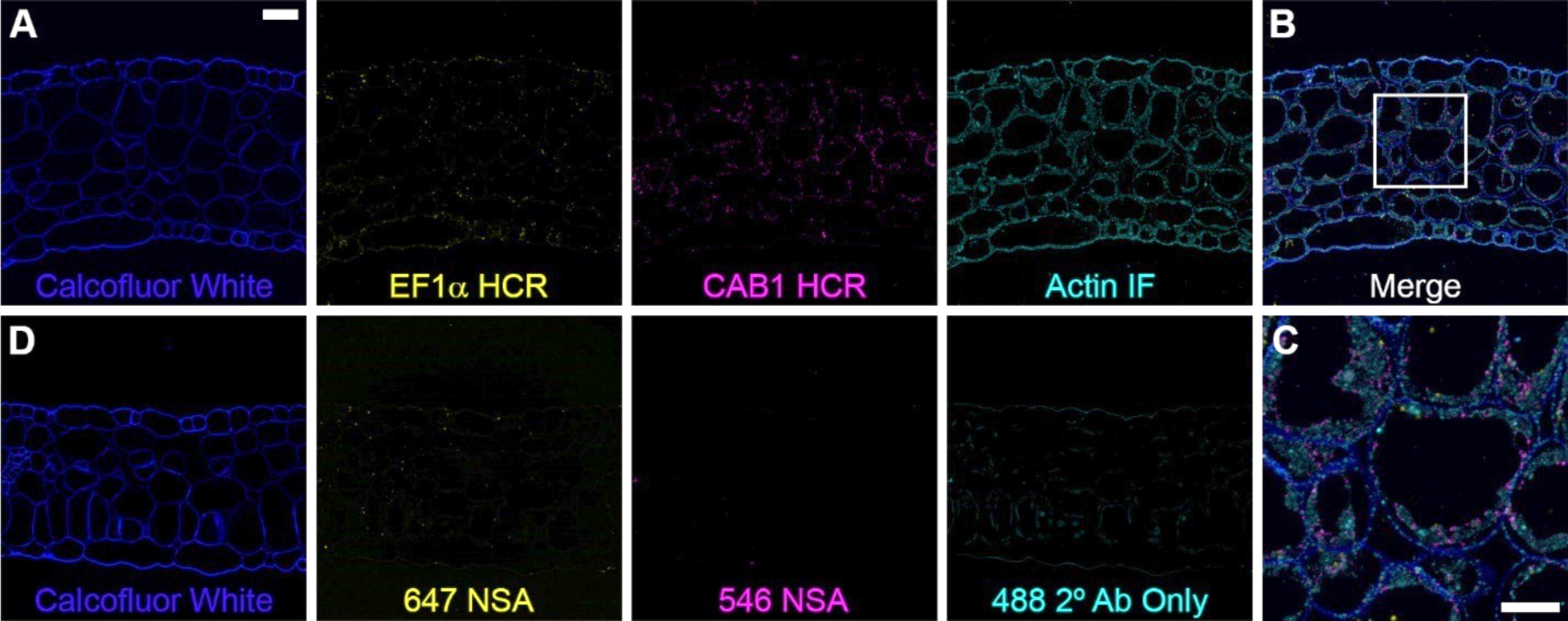
HCR can be used to detect mRNA transcripts and protein simultaneously in 0.5 µm methacrylate sections. Longitudinal plane through a 3-day *Arabidopsis thaliana* cotyledon embedded in methacrylate, sectioned at 0.5µm and imaged with a 63x/1.2 NA water immersion objective. A. Multiplex HCR with IHC. Left to right: Calcofluor White cell wall stain. HCR for EF1*α* (Alexa Fluor 647, Elongation factor 1 alpha). HCR for CAB1 (Alexa Fluor 546, Chlorophyll binding protein a/b). IHC for pan-Actin (DyLight 488). Scale = 20 µm. B. Merged image. C. Optical zoom of area from bounding box in B. Scale = 10 µm. D. Non-specific amplification negative control imaged as in A.

**Figure S5.**
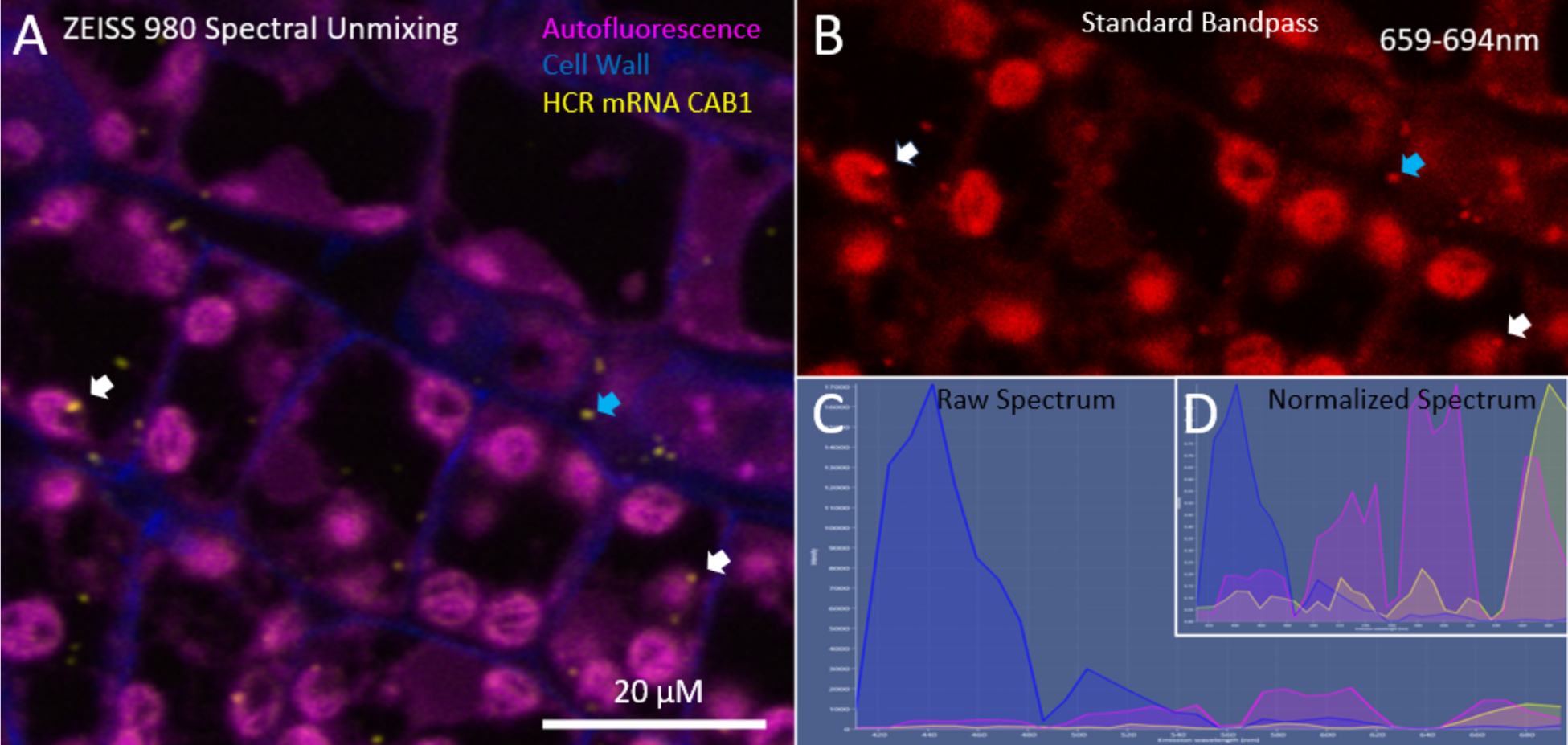
Linear spectral unmixing and spectra. A. Linear spectral unmixing of *A. thaliana* meristem to delineate the cell wall with Calcofluor White (blue), plant autofluorescence (magenta) and HCR of Alexa Fluor 650 labeled *CAB1* mRNA (yellow). B. Combined signal from the 659-694 nm (9 nm windows) channels mimicking a standard bandpass filter for Alexa Fluor 650 illustrated the extent of autofluorescence overlap an interference with *CAB1* mRNA probe (white arrows) while it was possible to distinguish some foci in some areas (blue arrows). This demonstrated the challenge to clearly localize and/or quantify some signals with highly autofluorescent tissue or weaker probe signals. (C) Raw and (D) Normalized spectrum of Calcofluor White (blue), Autofluorescence (magenta) and Alexa Fluor 650 (yellow) spectrum used for linear unmixing of (A) and showed the extent of autofluorescence overlap and dominated the green and orange/red emission region of the spectrum. Scale = 20 μm.

**Figure S6.**
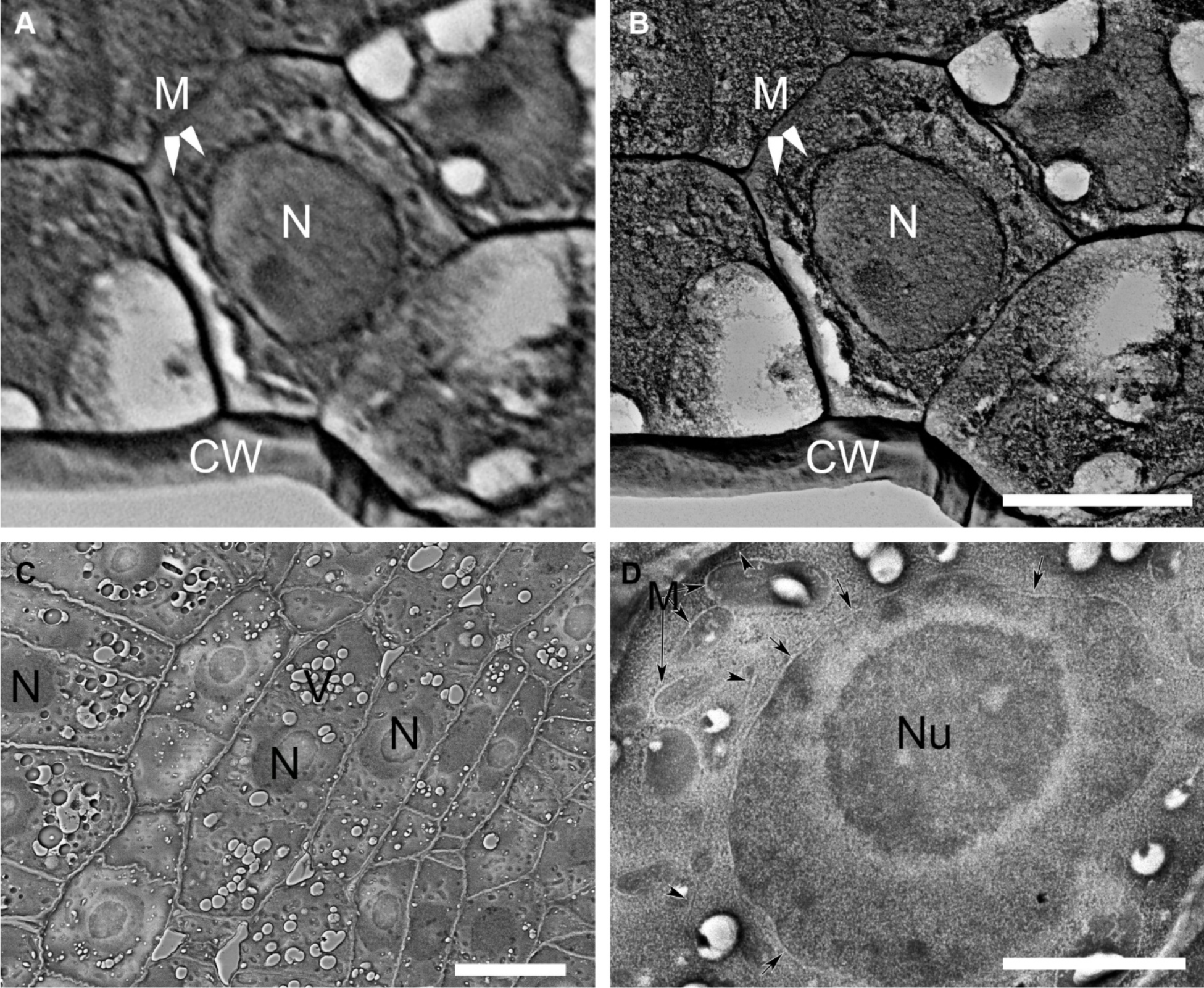
FESEM of methacrylate de-embedded plant tissues. A. *A thaliana* shoot meristem from 1.5 μm section (same specimen/adjacent sections as Fig 6) imaged using back-scatter electrons at 50nm pixel resolution and B the same location at 5 nm pixel resolution which showed improved delineation of cell organelles such as the nucleus (N), cell wall (CW) and mitochondria (M). Scale = 5 µm. C. A portion of an *A. thaliana* root meristem (0.5 μm thick) in cross-section acquired at 5nm pixel resolution, and at greater magnification (D) showed that nuclei (N), vacuoles (V), mitochondria (M), the nucleolus (Nu), nuclear envelope (arrows) and endoplasmic reticulum (arrow heads) are all readily discernable. Scale = 2.5 μm.

**Table S1.**
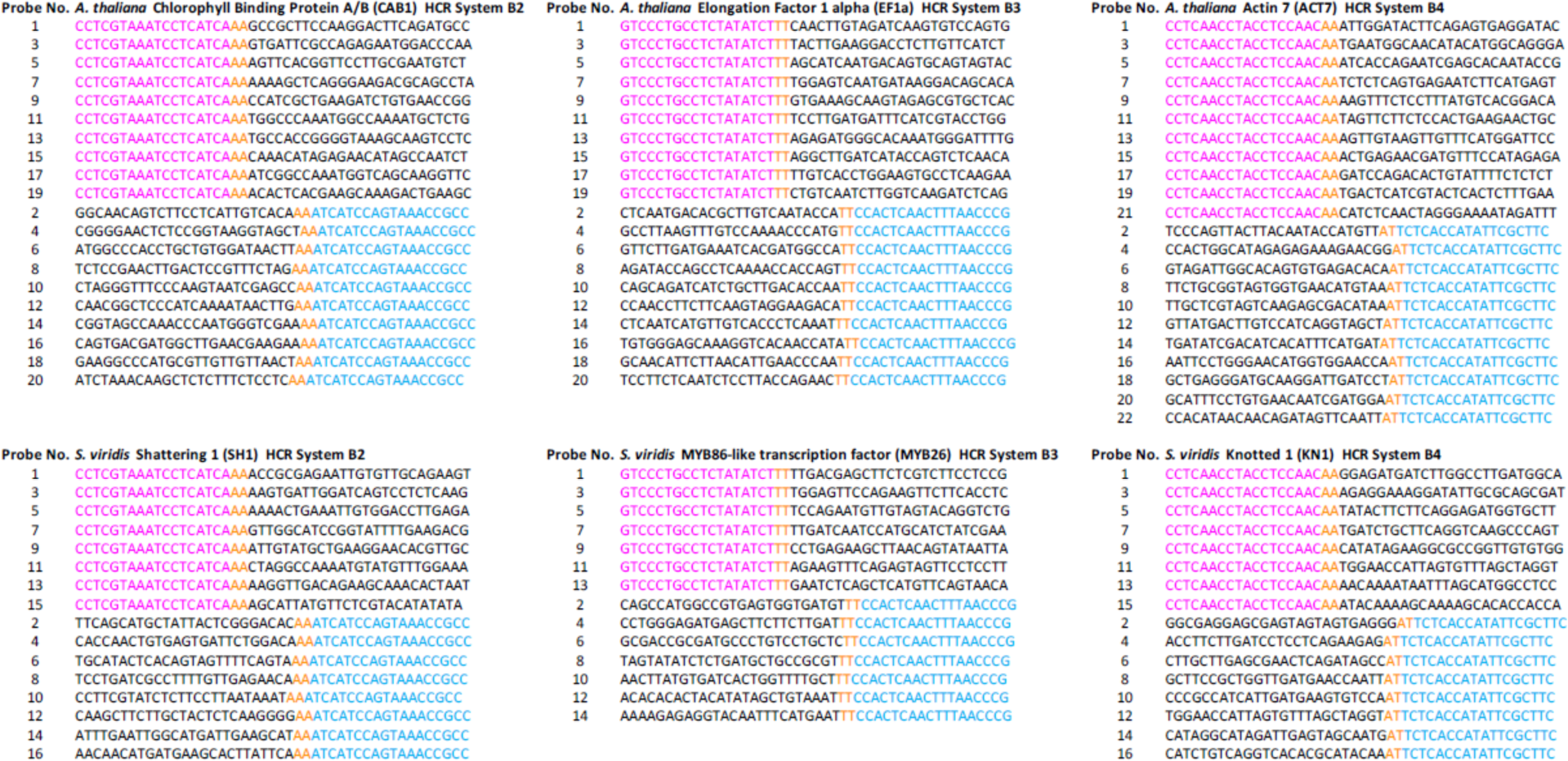
Sequences of HCR probes. All sequences are listed 5’ to 3’. The 5’ half of the initiator sequence is shown in magenta while the 3’ half of the initiator sequence is shown in cyan. The spacer sequences are shown in orange. Please refer to the Materials and Methods section for information on gene accession numbers and probe design.

## REFERENCES

1. Young AP, Jackson DJ, Wyeth RC. A technical review and guide to RNA fluorescence in situ hybridization. PeerJ [Internet]. 2020 Mar 19;8:e8806. Available from: https://peerj.com/articles/8806

2. Levsky JM, Singer RH. Fluorescence in situ hybridization: past, present and future. J Cell Sci [Internet]. 2003 Jul 15;116(14):2833–8. Available from: https://journals.biologists.com/jcs/article/116/14/2833/27272/Fluorescence-in-situ-hybridization-past-present

3. Jensen E. Technical Review: In Situ Hybridization. Anat Rec [Internet]. 2014 Aug 9;297(8):1349–53. Available from: https://anatomypubs.onlinelibrary.wiley.com/doi/10.1002/ar.22944

4. Hejátko J, Blilou I, Brewer PB, Friml J, Scheres B, Benková E. In situ hybridization technique for mRNA detection in whole mount Arabidopsis samples. Nat Protoc [Internet]. 2006 Nov 22;1(4):1939–46. Available from: https://www.nature.com/articles/nprot.2006.333

5. Kronenberger J, Desprez T, Höfte H, Caboche M, Traas J. A methacrylate embedding procedure developed for immunolocalization on plant tissues is also compatible with in situ hybridization. Cell Biol Int [Internet]. 1993 Nov 2;17(11):1013–21. Available from: https://onlinelibrary.wiley.com/doi/10.1006/cbir.1993.1031

6. Stahl Y, Simon R. mRNA Detection by Whole Mount In Situ Hybridization (WISH) or Sectioned Tissue In Situ Hybridization (SISH) in Arabidopsis. In 2010. p. 239–51. Available from: http://link.springer.com/10.1007/978-1-60761-765-5_16

7. Javelle M, Marco CF, Timmermans M. In Situ Hybridization for the Precise Localization of Transcripts in Plants. J Vis Exp [Internet]. 2011 Nov 23;(57). Available from: http://www.jove.com/details.php?id=3328

8. Rozier F, Mirabet V, Vernoux T, Das P. Analysis of 3D gene expression patterns in plants using whole-mount RNA in situ hybridization. Nat Protoc [Internet]. 2014 Oct 25;9(10):2464–75. Available from: https://www.nature.com/articles/nprot.2014.162

9. Zöllner NR, Bezrutczyk M, Laureyns R, Nelissen H, Simon R, Frommer WB. An RNA in situ hybridization protocol optimized for monocot tissue. STAR Protoc [Internet]. 2021 Jun;2(2):100398. Available from: https://linkinghub.elsevier.com/retrieve/pii/S2666166721001052

10. Raj A, van den Bogaard P, Rifkin SA, van Oudenaarden A, Tyagi S. Imaging individual mRNA molecules using multiple singly labeled probes. Nat Methods [Internet]. 2008 Oct 21;5(10):877–9. Available from: https://www.nature.com/articles/nmeth.1253

11. Duncan S, Olsson TSG, Hartley M, Dean C, Rosa S. A method for detecting single mRNA molecules in Arabidopsis thaliana. Plant Methods [Internet]. 2016 Dec 5;12(1):13. Available from: http://plantmethods.biomedcentral.com/articles/10.1186/s13007-016-0114-x

12. Yang W, Schuster C, Prunet N, Dong Q, Landrein B, Wightman R, et al. Visualization of Protein Coding, Long Noncoding, and Nuclear RNAs by Fluorescence in Situ Hybridization in Sections of Shoot Apical Meristems and Developing Flowers. Plant Physiol [Internet]. 2020 Jan;182(1):147–58. Available from: https://academic.oup.com/plphys/article/182/1/147-158/6116070

13. Nobori T, Oliva M, Lister R, Ecker JR. Multiplexed single-cell 3D spatial gene expression analysis in plant tissue using PHYTOMap. Nat Plants [Internet]. 2023 Jun 12;9(7):1026–33. Available from: https://www.nature.com/articles/s41477-023-01439-4

14. Wang F, Flanagan J, Su N, Wang L-C, Bui S, Nielson A, et al. RNAscope. J Mol Diagnostics [Internet]. 2012 Jan;14(1):22–9. Available from: https://linkinghub.elsevier.com/retrieve/pii/S1525157811002571

15. Choi HMT, Beck VA, Pierce NA. Next-Generation in Situ Hybridization Chain Reaction: Higher Gain, Lower Cost, Greater Durability. ACS Nano [Internet]. 2014 May 27;8(5):4284–94. Available from: https://pubs.acs.org/doi/10.1021/nn405717p

16. Choi HMT, Schwarzkopf M, Fornace ME, Acharya A, Artavanis G, Stegmaier J, et al. Third-generation in situ hybridization chain reaction: Multiplexed, quantitative, sensitive, versatile, robust. Dev. 2018;145(12):1–10.

17. Speel EJM, Hopman AHN, Komminoth P. Tyramide Signal Amplification for DNA and mRNA In Situ Hybridization. In: In Situ Hybridization Protocols [Internet]. New Jersey: Humana Press; p. 33–60. Available from: http://link.springer.com/10.1385/1-59745-007-3:33

18. Ke R, Mignardi M, Pacureanu A, Svedlund J, Botling J, Wählby C, et al. In situ sequencing for RNA analysis in preserved tissue and cells. Nat Methods [Internet]. 2013 Sep 14;10(9):857–60. Available from: https://www.nature.com/articles/nmeth.2563

19. Gyllborg D, Langseth CM, Qian X, Choi E, Salas SM, Hilscher MM, et al. Hybridization-based in situ sequencing (HybISS) for spatially resolved transcriptomics in human and mouse brain tissue. Nucleic Acids Res [Internet]. 2020 Nov 4;48(19):e112–e112. Available from: https://academic.oup.com/nar/article/48/19/e112/5912821

20. Bowling AJ, Pence HE, Church JB. Application of a novel and automated branched DNA in situ hybridization method for the rapid and sensitive localization of mRNA molecules in plant tissues. Appl Plant Sci [Internet]. 2014 Apr 3;2(4). Available from: https://bsapubs.onlinelibrary.wiley.com/doi/10.3732/apps.1400011

21. Munganyinka E, Margaria P, Sheat S, Ateka EM, Tairo F, Ndunguru J, et al. Localization of cassava brown streak virus in Nicotiana rustica and cassava Manihot esculenta (Crantz) using RNAscope® in situ hybridization. Virol J [Internet]. 2018 Dec 14;15(1):128. Available from: https://virologyj.biomedcentral.com/articles/10.1186/s12985-018-1038-z

22. Solanki S, Ameen G, Zhao J, Flaten J, Borowicz P, Brueggeman RS. Visualization of spatial gene expression in plants by modified RNAscope fluorescent in situ hybridization. Plant Methods [Internet]. 2020 Dec 15;16(1):71. Available from: https://plantmethods.biomedcentral.com/articles/10.1186/s13007-020-00614-4

23. Choi HMT, Schwarzkopf M, Fornace ME, Acharya A, Artavanis G, Stegmaier J, et al. Third-generation in situ hybridization chain reaction: multiplexed, quantitative, sensitive, versatile, robust. Development [Internet]. 2018 Jun 15;145(12). Available from: https://journals.biologists.com/dev/article/145/12/dev165753/48466/Third-generation-in-situ-hybridization-chain

24. Trivedi V, Choi HMT, Fraser SE, Pierce NA. Multidimensional quantitative analysis of mRNA expression within intact vertebrate embryos. Development [Internet]. 2018 Jan 1;145(1). Available from: https://journals.biologists.com/dev/article/145/1/dev156869/48765/Multidimensional-quantitative-analysis-of-mRNA

25. Huang T, Guillotin B, Rahni R, Birnbaum KD, Wagner D. A rapid and sensitive, multiplex, whole mount RNA fluorescence in situ hybridization and immunohistochemistry protocol. Plant Methods [Internet]. 2023 Nov 22;19(1):131. Available from: https://plantmethods.biomedcentral.com/articles/10.1186/s13007-023-01108-9

26. Oliva M, Stuart T, Tang D, Pflueger J, Poppe D, Jabbari JS, et al. An environmentally responsive transcriptional state modulates cell identities during root development. bioRxiv [Internet]. 2022;2022.03.04.483008. Available from: https://www.biorxiv.org/content/10.1101/2022.03.04.483008v2%0A https://www.biorxiv.org/content/10.1101/2022.03.04.483008v2.abstract

27. Lherminier J, Bonfiglioli RG, Daire X, Symons RH, Boudon-Padieu E. Oligodeoxynucleotides as probes forin situhybridization with transmission electron microscopy to specifically localize phytoplasma in plant cells. Mol Cell Probes [Internet]. 1999 Feb;13(1):41–7. Available from: https://linkinghub.elsevier.com/retrieve/pii/S0890850898902134

28. Motte PM, Loppes R, Menager M, Deltour R. Three-dimensional electron microscopy of ribosomal chromatin in two higher plants: a cytochemical, immunocytochemical, and in situ hybridization approach. J Histochem Cytochem [Internet]. 1991 Nov 5;39(11):1495–506. Available from: http://journals.sagepub.com/doi/10.1177/39.11.1918926

29. Harris N, Croy RRD. Localization of mRNA for pea legumin:In situ hybridization using a biotinylated cDNA probe. Protoplasma [Internet]. 1986 Feb;130(1):57–67. Available from: http://link.springer.com/10.1007/BF01283331

30. Warren KC, Coyne KJ, Waite JH, Cary SC. Use of Methacrylate De-embedding Protocols for In Situ Hybridization on Semithin Plastic Sections with Multiple Detection Strategies. J Histochem Cytochem [Internet]. 1998 Feb 26;46(2):149–55. Available from: http://journals.sagepub.com/doi/10.1177/002215549804600203

31. Mueller M, Wacker K, Hickey WF, Ringelstein EB, Kiefer R. Co-Localization of Multiple Antigens and Specific DNA. Am J Pathol [Internet]. 2000 Dec;157(6):1829–38. Available from: https://linkinghub.elsevier.com/retrieve/pii/S0002944010648225

32. Yu Y, Hu H, Doust AN, Kellogg EA. Divergent gene expression networks underlie morphological diversity of abscission zones in grasses. New Phytol [Internet]. 2020 Feb 28;225(4):1799–815. Available from: https://nph.onlinelibrary.wiley.com/doi/10.1111/nph.16087

33. Jackson D, Veit B, Hake S. Expression of maize KNOTTED1 related homeobox genes in the shoot apical meristem predicts patterns of morphogenesis in the vegetative shoot. Development [Internet]. 1994 Feb 1;120(2):405–13. Available from: https://journals.biologists.com/dev/article/120/2/405/38273/Expression-of-maize-KNOTTED1-related-homeobox

34. Bolduc N, Yilmaz A, Mejia-Guerra MK, Morohashi K, O’Connor D, Grotewold E, et al. Unraveling the KNOTTED1 regulatory network in maize meristems. Genes Dev [Internet]. 2012 Aug 1;26(15):1685–90. Available from: http://genesdev.cshlp.org/lookup/doi/10.1101/gad.193433.112

35. Brewer PB, Heisler MG, Hejátko J, Friml J, Benková E. In situ hybridization for mRNA detection in Arabidopsis tissue sections. Nat Protoc [Internet]. 2006 Aug 9;1(3):1462–7. Available from: https://www.nature.com/articles/nprot.2006.226

36. Donaldson L. Autofluorescence in Plants. Molecules [Internet]. 2020 May 21;25(10):2393. Available from: https://www.mdpi.com/1420-3049/25/10/2393

37. De Mazière A, van der Beek J, van Dijk S, de Heus C, Reggiori F, Koike M, et al. An optimized protocol for immuno-electron microscopy of endogenous LC3. Autophagy [Internet]. 2022 Dec 2;18(12):3004–22. Available from: https://www.tandfonline.com/doi/full/10.1080/15548627.2022.2056864

38. Mansfield JR, Gossage KW, Hoyt CC, Levenson RM. Autofluorescence removal, multiplexing, and automated analysis methods for in-vivo fluorescence imaging. J Biomed Opt [Internet]. 2005;10(4):041207. Available from: http://biomedicaloptics.spiedigitallibrary.org/article.aspx?doi=10.1117/1.2032458

39. Huang K, Demirci F, Batish M, Treible W, Meyers BC, Caplan JL. Quantitative, super-resolution localization of small RNAs with sRNA-PAINT. Nucleic Acids Res. 2020;48(16):e96.

40. McDowell JM, An Y, Huang S, McKinney EC, Meagher RB. The Arabidopsis ACT7 Actin Gene Is Expressed in Rapidly Developing Tissues and Responds to Several External Stimuli. Plant Physiol [Internet]. 1996 Jul 1;111(3):699–711. Available from: https://academic.oup.com/plphys/article/111/3/699/6070322

41. Huang K, Batish M, Teng C, Harkess A, Meyers BC, Caplan JL. Quantitative Fluorescence In Situ Hybridization Detection of Plant mRNAs with Single-Molecule Resolution. In 2020. p. 23–33. Available from: http://link.springer.com/10.1007/978-1-0716-0712-1_2

42. Huang G, Eckrich S. Quantitative Fluorescent in situ Hybridization Reveals Differential Transcription Profile Sharpening of Endocytic Proteins in Cochlear Hair Cells Upon Maturation. Front Cell Neurosci [Internet]. 2021 Feb 26;15. Available from: https://www.frontiersin.org/articles/10.3389/fncel.2021.643517/full

43. No Title [Internet]. Available from: https://www.biosearchtech.com/support/tools/design-software/stellaris-probe-designer

44. Schindelin J, Arganda-Carreras I, Frise E, Kaynig V, Longair M, Pietzsch T, et al. Fiji: an open-source platform for biological-image analysis. Nat Methods [Internet]. 2012 Jul 28;9(7):676–82. Available from: http://www.nature.com/articles/nmeth.2019

45. Collman F, Buchanan J, Phend KD, Micheva KD, Weinberg RJ, Smith SJ. Mapping Synapses by Conjugate Light-Electron Array Tomography. J Neurosci [Internet]. 2015 Apr 8;35(14):5792–807. Available from: https://www.jneurosci.org/lookup/doi/10.1523/JNEUROSCI.4274-14.2015

46. A Stable Lead by Modification of Sato’s Method. J Electron Microsc (Tokyo) [Internet]. 1986 Oct; Available from: https://academic.oup.com/jmicro/article/35/3/304/847425/A-Stable-Lead-by-Modification-of-Satos-Method

